# Pharmacological inhibition of CXCR4 increases the anti-tumor activity of conventional and targeted therapies in B-cell lymphoma models

**DOI:** 10.1101/2025.07.10.664047

**Authors:** Laura Barnabei, Alberto J. Arribas, Luciano Cascione, Francesca Guidetti, Filippo Spriano, Chiara Tarantelli, Samuele Di Cristofano, Stefano Raniolo, Andrea Rinaldi, Anastasios Stathis, Davide Rossi, Emanuele Zucca, Vittorio Limongelli, Johann Zimmermann, Francesco Bertoni

## Abstract

**Background:** CXCR4 is a chemokine receptor frequently implicated in the pathogenesis and treatment resistance of B-cell lymphomas and other tumor types. Thus, CXCR4 targeting is explored using various approaches, including small molecules, antibodies, and short peptides. Here, we investigated the antitumor effects of the CXCR4 pharmacological inhibition with the synthetic peptides SPX5551 and balixafortide, as single agents and in combination, in various B-cell lymphoma models.

**Methods:** Binding modalities were assessed via molecular docking and simulation. In vitro assays evaluated single-agent and combinatorial effects of CXCR4 antagonists with BTK, PI3K, and conventional therapies across 20 lymphoma cell lines, including models with acquired resistance. Transcriptomic analyses and mechanistic studies elucidated the pathways modulated by combined CXCR4 and BTK inhibition.

**Results:** Structural modeling confirmed similar CXCR4 binding modes for SPX5551 and balixafortide. SPX5551 displayed minimal single-agent activity but restored sensitivity to BTK and PI3K inhibitors in resistant marginal zone lymphoma (MZL) models. Across mantle cell lymphoma (MCL), chronic lymphocytic leukemia (CLL), and diffuse large B-cell lymphoma (DLBCL) models, SPX5551 enhanced the efficacy and/or potency of ibrutinib, copanlisib, rituximab, and R-CHOP. In MCL, co-treatment with SPX5551 and ibrutinib synergistically induced apoptosis, inhibited NF-κB and AKT signaling, and led to broader transcriptomic repression of tumor-promoting pathways compared to either agent alone.

**Conclusions:** Although CXCR4 inhibitors alone show limited cytotoxicity, their combination with standard and targeted therapies significantly enhances anti-lymphoma effects, particularly in drug-resistant settings. These findings provide a strong rationale for clinical evaluation of CXCR4 blockade as a combinatorial strategy in B-cell lymphomas.

## Introduction

C-X-C chemokine receptor type 4 (CXCR4) is a chemokine receptor belonging to the family of the rhodopsin-like G protein-coupled receptors, ubiquitously expressed in various normal and neoplastic cell types, including B-cell lymphoid neoplasms ^1,2^. The CXCL12, also known as SDF1α, is the canonical ligand of CXCR4 and, once binds, initiates a wide network of downstream signal transduction. The main signaling pathways include JAK/STAT, PI3K/AKT/mTOR, RAS/RAF, MAPK, and NF-κB pathways, contributing to tumor angiogenesis, metastasis, cell survival, and immune evasion. The gene coding CXCR4 is also recurrently mutated in human cancers, especially in Waldenstrom macroglobulinemia (WM)/IgM lymphoplasmacytic lymphoma (LPL). *CXCR4* somatic mutations, detected in up to 40% of WM patients, resemble the germline mutations detected in patients with Warts, hypogammaglobulinemia, infections, and myelokathexis (WHIM) syndrome ^3^. The mutations largely occur in the C-terminal domain of CXCR4, determining the loss of regulatory serines, which are phosphorylated after CXCL12-induced CXCR4 activation and lead to constitutive active downstream signaling ^3^.

Based on this evidence, CXCR4 is an attractive molecular target for developing anticancer therapeutics and targeted radioimaging ^1,2,4^. CXCR4 targeting is explored using various approaches, including small molecules, antibodies, and short peptides ^1,2,4,5^. Two CXCR4 antagonists, plerixafor (AMD3100) and motixafortide (BL-8040; BKT-140, 4F-benzoyl-TN14003, peptide T140), are approved by the US Food and Drug Administration (FDA) for hematopoietic stem cell mobilization for the preparation of autologous stem cell transplantations in cancer patients, and an additional one, mavorixafor (X4P-001, AMD11070, AMD070), for WHIM syndrome patients ^6^. SPX5551 (POL5551) and its close analog balixafortide (SPX6326, POL6326) are synthetic peptides obtained via the protein epitope mimetic (PEM) approach following the optimization of libraries initially designed starting from polyphemusin II, an 18-amino acid peptide isolated from *Limulus polyphemus* (the Atlantic horseshoe crab) and its close analog T22 ([Tyr5, 12, Lys7]-polyphemusin II) ^5^. Balixafortide has entered the clinical evaluation as a hematopoietic stem cell mobilizer ^7^ and in combination with the tubulin-targeting agent eribulin in breast cancer patients ^8^. Preclinical evidence has shown the potential benefit of combining SPX5551 and balixafortide with anti-cancer compounds in solid tumors or acute leukemia models ^9–11^. Importantly, in lymphoid neoplasms, CXCR4 levels are upregulated by B cell receptor (BCR)/PI3K blockade ^12,13^, and high CXCR4 expression is associated with resistance to BCR/PI3K inhibitors ^13,14^, and the presence of *CXCR4* gene mutations might reduce the sensitivity to BTK inhibitors in WM patients ^15^.

Here, we explored the pharmacological inhibition of CXCR4 using SPX5551 and balixafortide in lymphoma models as a single agent and in combination with drugs used in the clinics.

## Materials and Methods

### Computational Studies

We studied the binding of balixafortide and SPX5551 to human CXCR4 (UniProt P61073-1 sequence, residues 25-303) employing the crystallographic atomistic structure of the CXCR4 receptor in complex with the CVX15 cyclic peptide (PDBID 3OE0) ^16^, a peptide structurally similar to SPX5551 and balixafortide. Missing residues in the experimental structure, comprising intracellular loops (residues 67-70 and 229-238) were modelled using the model generator and loop optimization algorithm of Modeller ^17,18^. Balixafortide and SPX5551 were obtained by manually mutating the CVX15 cyclic peptide bound to CXCR4 in the X-ray structure 3OE0 using ChimeraX 1.9 ^19^. Although the high identity among the three cyclic peptides suggests a similarity in mechanism, we explored potential alternative binding modes of balixafortide and SPX5551 bound to the orthosteric site of the CXCR4 receptor. To do so, we performed rigid body perturbations through the CycpepRigidBodyPermutationMover in the “randomized_perturbation” mode, as implemented in Rosetta (v3.14) ^20^. A random offset of 0.8 Å (flag “*random_position_offset*”) and 2 degrees (flag “*random_orientation_perturbation*”) was applied to the peptide position and rotation axis in each permutation cycle, respectively. Given the good structural symmetry of our peptides, we allowed permutations including those that align the peptide to a reversed permutation of itself. After every rigid body perturbation, side-chain repacking and all-atom energy minimization were conducted using the FastRelax protocol in Rosetta. At the end of each cycle, we determined the total score of each model using the ref2015 Rosetta energy function (REU) ^21^. This cycle was repeated 5000 times, generating a total of 10000 docked conformations of balixafortide and SPX5551 together. Finally, we selected the peptide binding mode with the best REU score. The poses found by the docking calculations were analyzed by plotting the peptide root mean square deviation (RMSD) vs. the REU score (Supplementary Figure S1). The peptide RMSD values were calculated for the Cα atoms of balixafortide and SPX5551 in each docking-derived conformation relative to the top-scoring REU pose, following alignment of the Cα atoms of the CXCR4 binding pocket residues. As shown in Supplementary Figure S1, higher RMSD values correspond to poses with poor docking scores, while lower RMSD values indicate better-scoring conformations, highlighting multiple peptide binding modes closely resembling the top scored pose.

The selected receptor/peptide complexes were then embedded in a 105 × 105 A° (along x and y axes) pre-equilibrated POPC/cholesterol (7:3 molar ratio) bilayer and solvated using the TIP3P water model with the aid of the membrane-builder tool of CHARMM-GUI.org (http://www.charmm-gui.org), using the CHARMM36m forcefield for protein and lipids ^22,23^. The GROMACS 2021.5 ^24^ code was used to perform all-atom energy minimization in explicit solvent. A cutoff of 12 Å was used for short-range interactions. The long-range electrostatic interactions were computed through the particle mesh Ewald (PME) method ^25^ using a 1.0 Å grid spacing in periodic boundary conditions. The non-iterative LINCS ^26^ algorithm was applied to constrain bonds. To solve all the steric clashes, each system underwent 7 consecutive phases of steepest descent energy minimization, gradually releasing position restraints from an initial force constant of 4000 kJ/mol/nm^2^ for backbone atoms, 2000 kJ/mol/nm^2^ for side-chain atoms, and 1000 kJ/mol/nm^2^ for lipid atoms (phase 1) to 0 kJ/mol/nm^2^ for the whole system (phases 6 and 7). Each minimization phase was stopped when the maximum force reached was less than 100.0 kJ/mol/nm^2^. Figures of the final binding mode of balixafortide and SPX5551 to CXCR4 in explicit solvent were generated using ChimeraX 1.9 ^19^.

### Cell Lines

Lymphoma cell lines (Supplementary Table S1) were cultured according to the recommended conditions. All media were supplemented with fetal bovine serum (FBS) (10% or 20%) and penicillin-streptomycin-neomycin (≈5,000 units penicillin, 5 mg streptomycin, and 10 mg neomycin/mL; Sigma-Aldrich, Darmstadt, Germany). Cell line identities were confirmed by short tandem repeat DNA fingerprinting using the Promega GenePrint 10 System kit (B9510). Cells were periodically tested for mycoplasma negativity using the MycoAlert Mycoplasma Detection Kit (Lonza, Visp, Switzerland).

### Compounds

SPX5551 and balixafortide were provided by Spexis. Ibrutinib, copanlisib, idelalisib, doxorubicin, vincristine, and prednisolone were purchased from Selleckchem (Houston, TX, USA). Rituximab was purchased from Roche (Basel, Switzerland), and 4-hydroperoxy-cyclophosphamide from Santa Cruz Biotechnology (Heidelberg, Germany).

### In vitro cytotoxic activity

Cells were manually seeded in 96-well plates at their optimal cell concentration (Supplementary Table S2). Treatments were performed manually. After 72 hours, cell viability was determined by using 3-(4,5-dimethyl-thiazol-2-yl)-2,5-diphenyltetrazolium bromide (MTT), and the reaction was stopped after 4 hours with sodium dodecyl sulfate lysis buffer.

For combination studies, cells were exposed (72 hours) to five increasing concentrations of the two agents, either alone or in combination, followed by an MTT assay. The concentrations used for all treatments are mentioned in Supplementary Table S2. For R-CHOP treatment, cells were exposed for 72 h to 0.1 μg/mL CHOP + 100 μg/mL rituximab to five different concentrations in serial dilution 1:10. Rituximab was diluted to clinically recommended serum levels ^27^ and CHOP represented a mix reflecting the clinical ratios of the drugs (85%, 4-hydroperoxy-cyclophosphamide; 5.5%, doxorubicin; 0.16%, vincristine; 11.1%, prednisolone) ^28^.

Sensitivity to single drug treatments was evaluated by IC50 (4-parameter calculation upon log-scaled doses) and area under the curve (PharmacoGX R package ^29^) calculations. The beneficial effect of the combinations versus the single agents was considered both as synergism according to the Chou-Talalay combination index ^30^, and as efficacy (maximal effect of the treatment) and potency (defined as minimal dose with a biological effect) according to the MuSyC algorithm ^31^. Three independent experiments were performed for each combination.

### Flow Cytometry

Cells were washed in cold PBS and fixed in cold 70% ethanol overnight. Cells were then stained with PI (50 mg/mL; Sigma-Aldrich) for cell cycle, or double-stained with Annexin V–FITC (Sigma-Aldrich)/PI, for apoptosis. Surface expression of CXCR4 was determined with the CXCR4-PE (1:200 dilution; BD Bioscience #561733). Cells were analyzed with a FACS Canto II instrument (BD Biosciences). Each sample’s median fluorescence intensity (MFI) was determined using FACSDiva v8.0.1 software (BD Biosciences, Allschwil, Switzerland). Surface CXCR4 expression and the induction of apoptosis and distribution of cells in distinct cell cycle phases following the treatment were evaluated with the flowCore R package ^32^.

### Immunofluorescence and confocal microscopy

After exposure to the compounds of interest, coating of the cells was done on a poly-L-lysine matrix. The cells were then subjected to fixation using 4% paraformaldehyde (PFA) for 20 min at room temperature. The PBS+ 0.1% Triton X-100 treatment for 10 mins at RT was used for the permeabilization of cells. 5% BSA (TBST) for 1 h was used to block any unspecific staining prior to the staining process. The 5% bovine serum albumin (BSA) (TBST) was used to dilute the antibodies. It was followed by the incubation of samples at 4°C with primary rabbit monoclonal anti-human NF-kB p65 antibody (D14E12) (1/100; Cell Signaling) overnight. Subsequently, the cells were incubated with the Alexa 568 labelled secondary goat antibody anti-rabbit IgG (Thermo Fisher Scientific) for an hour at RT in the dark. The counterstain was done with 0.3 g/mL of 4,6-diamidino-2-phenylindole Sigma-Aldrich, Buchs, Switzerland), after washing the cells three times with buffered saline. The images were rendered on a Leica SP5 (Heerbrugg, Switzerland) equipped with an objective lens with magnification. The protein quantification was gauged by ImageJ software ^33^.

### RNA sequencing and data mining

Total RNA was extracted using the All Prep DNA/RNA Mini Kit (Qiagen, Hilden, Germany) according to the manufacturer’s instructions. RNA concentration and integrity were measured with a NanoDrop ND-1000 spectrophotometer and assessed through the Bioanalyzer 2100 instrument (Agilent Technologies, Santa Clara, CA, USA). NEBNext Ultra Directional RNA Library Prep Kit for Illumina (New England BioLabs Inc.) was employed with the NEBNext Multiplex Oligos for Illumina (New England BioLabs Inc.) and NEBNext Poly(A) mRNA Magnetic Isolation Module for cDNA synthesis with the addition of barcode sequences. The pre-pool sequencing was performed using the NextSeq2000 (Illumina, San Diego, CA, USA) with the P2 reagents kit V3 (100 cycles; Illumina). Samples were processed starting from stranded, single-ended 120bp-long sequencing reads. Differentially expressed genes were calculated with a moderated t-test on RNA-seq. P-value <0.05 and absolute log-2 fold-change higher than 1 were considered significant. Functional analysis was performed on the collapsed gene symbol list using GSEA (Gene Set Enrichment Analysis) with the MSigDB (Molecular Signatures Database) C2-C7 gene sets ^34^, and the SignatureDB database (https://lymphochip.nih.gov/signaturedb/). Statistical tests were performed using the R environment (R Studio console; RStudio, Boston, MA, USA).

### *CXCR4* mutational status

*CXCR4* mutational status was obtained from previous publications containing whole-exome sequencing ^14,35,36^, from the COSMIC Cell Lines Project (v101 released 19-NOV-24) ^37^ or analyzing RNA-Seq data previously obtained in the cell lines ^38^. For mutation calling from RNA-Seq, sequencing reads were first aligned to the human reference genome (GRCh38) using STAR (v2.7.10a) in 2-pass mode to improve splice junction detection. Duplicates were marked using Picard, and base quality score recalibration was performed with GATK (v4.4.0) according to best practices for RNA-Seq. Variants were then called using GATK HaplotypeCaller in RNA-Seq mode, and resulting variant call format (VCF) files were filtered based on standard quality metrics (e.g., QD > 2.0, FS < 30.0, MQ > 40.0). Identified variants in CXCR4 were annotated using Ensembl VEP (Variant Effect Predictor) to assess their predicted functional impact. Only non-synonymous variants with moderate or high predicted impact were considered for further analysis. Variants were cross-referenced with known *CXCR4* mutations reported in the literature and COSMIC to distinguish somatic or recurrent alterations of potential biological relevance.

## Results

### SPX5551 and balixafortide share a similar binding mode to CXCR4

We investigated the binding modes of the two synthetic peptides SPX5551 and balixafortide to CXCR4 using docking calculations (Figure 1). As shown in Supplementary Figure S2, the predicted binding poses closely resemble the experimental binding conformation of CVX15, a cyclic peptide co-crystallized with CXCR4. Specifically, when aligning the C_α_ atoms of the CXCR4 binding pocket residues within 4 Å of the ligand peptides, the RMSD of SPX5551 and balixafortide’s C_α_ atoms relative to those of CVX15 is less than 1.5 Å, underscoring the structural similarity of their binding modes (Supplementary Figure S2). In both SPX5551 and balixafortide, Tyr1 might form a direct or water-mediated H-bond with Glu26 at the N-terminus of CXCR4, whereas His2 interacts with Phe189 of ECL2 through a π-π stacking interaction (Figure 1). The hydroxyl group of SPX5551’s Tyr3 H-bonds with His281, favouring the formation of an intramolecular T-shaped stacking interaction of this residue with Tyr12. In both SPX5551 and balixafortide, the hydroxyl group of Ser5 forms an H-bond with the Asp262’s carboxyl group of TM6, whereas the non-canonical DAB8 residue protrudes towards Asp97 (TM2) and Asp187 (ECL2), forming salt bridge interactions. Furthermore, Arg9 and Tyr10 H-bond with CXCR4’s Thr117 (TM3) and Tyr255 (TM6), whereas Lys14 engages a salt bridge interaction with Asp193 of TM5. It is interesting to note that the backbone atoms of Tyr10 and Cys11 form H-bonds with CXCR4’s backbone atoms of Arg188 and Tyr190, extending the β-strand of the ECL2 and further stabilizing the peptides’ binding mode.

**Figure 1.**
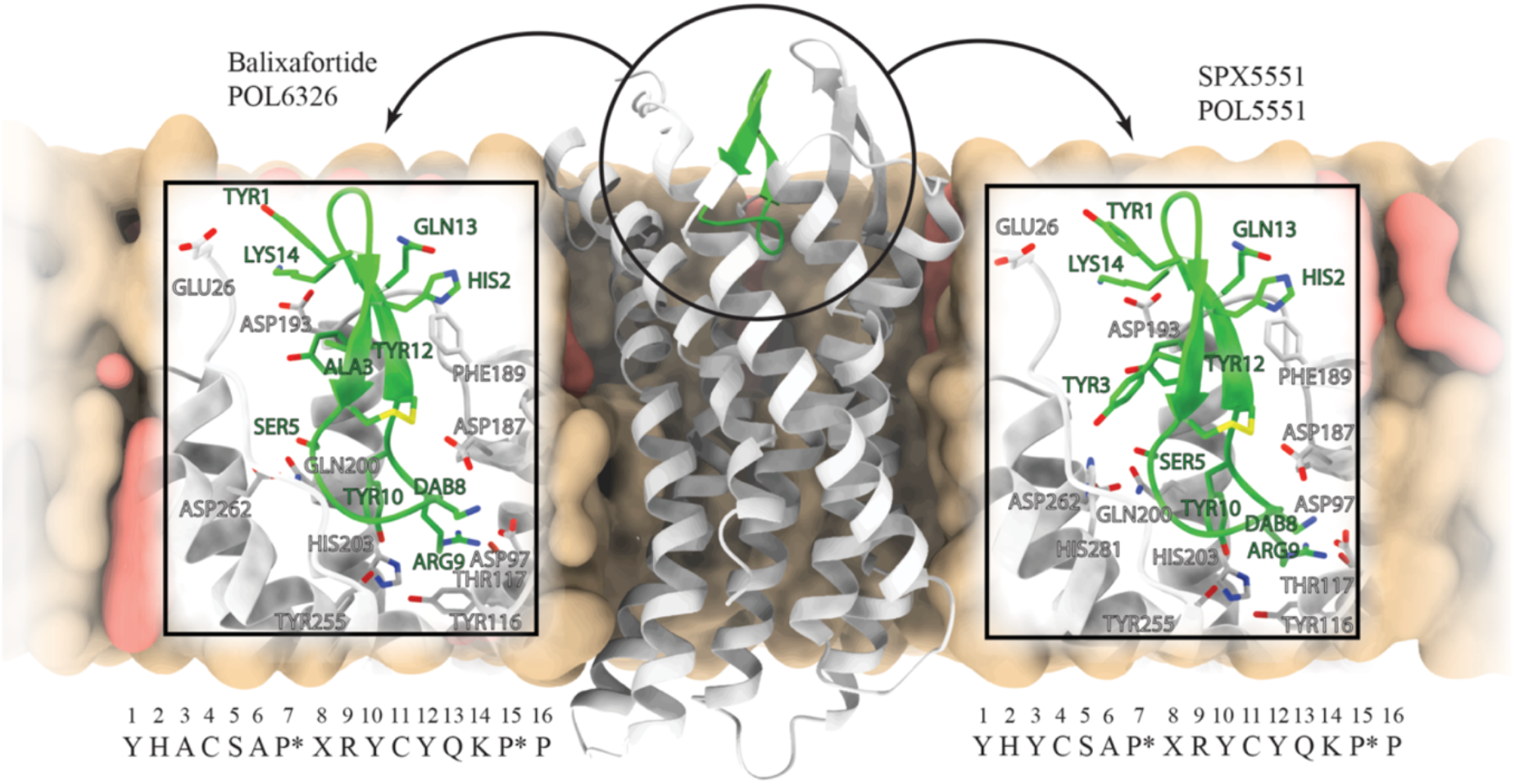
Docking of SPX5551 (POL5551) and balixafortide (SPX6326, POL6326) in CXCR4. Three-dimensional representation of the binding mode of balixafortide and SPX5551 to CXCR4 predicted by docking calculations. The protein is shown in grey ribbons, the cyclic peptides in green ribbons, and 1-Palmitoyl-2-oleoylphosphatidylcholine (POPC) and cholesterol are represented as surface in colors tan and salmon, respectively. The insets provide a magnified view of the binding modes for the two peptides. The sequence of each peptide is reported at the bottom of the corresponding figure as one-letter code. The two prolines in the D configuration are noted with an asterisk, whereas the 2-4-diaminobutyric acid is reported as X in the sequences and labeled as DAB in the insets. Peptide-CXCR4 interactions are shown as dashed black lines, whereas hydrogens are omitted for clarity.

### SPX5551 addition restores sensitivity to BTK and PI3K inhibitors in marginal zone lymphoma models of acquired secondary resistance

Given the binding mode similarity between SPX5551 and balixafortide, and since CXCR4 upregulation can contribute to resistance to BCR signaling inhibitors ^39,40^, we assessed the two peptides in two marginal zone lymphoma (MZL) cell lines, VL51 and Karpas1718, and in their derivatives with acquired resistance to BTK inhibitors and to PI3K inhibitors (copanlisib and idelalisib) obtained from ^14,35,36,41^ as a single agent and in combination.

After 72 hours of drug exposure, consistent with the high expression of surface CXCR4 in Karpas1718 cells, we observed a dose-dependent anti-proliferative activity only in the Karpas1718 models, stronger in the parental cell (IC50, 7.88 μM) than in its derivative with acquired resistance to BTK and PI3K inhibitors ^35^ (Supplementary Figure S3, Supplementary Figure S4, Supplementary Table S3). No effect was observed in the parental VL51 cells and their three derivatives with resistance to ibrutinib ^42^, idelalisib ^14^, or copanlisib ^41^ with any of the two compounds at concentrations up to 50 µM (Supplementary Figure S4), despite the expression of CXCR4 on the cells’ surface (Supplementary Figure S3) ^14^.

When we added the CXCR4 antagonist SPX5551 to the BTK inhibitor ibrutinib, the combination was strongly synergistic in VL51 and Karpas1718 parental cells and in their ibrutinib-resistant models (Supplementary Figure S5A): adding SPX5551 in a concentration ranging from 0.08 to 50 µM restored the sensitivity to ibrutinib in the ibrutinib-resistant cells (Supplementary Figure S5D). Although a dose-dependent decrease in cell viability was observed for the ibrutinib/SPX5551 combination in both cell lines, a strong anti-proliferative activity was observed in the Karpas1718 model.

Similarly, SPX5551 exhibited synergistic behaviour with the PI3Kδ inhibitor idelalisib and the PI3Kα/δ inhibitor copanlisib (Supplementary Figure S5B-C), restoring sensitivity to the PI3K inhibitors in resistant models (Supplementary Figure S5D). Of note, combinations with both BTK and PI3K inhibitors were more effective in the Karpas1718 models than those derived from VL51, consistent with the above-reported increased sensitivity to SPX5551 in the Karpas1718 than in the VL51 parental cells. The benefit of adding copanlisib was due to an improved efficacy rather than potency (Supplementary Figure S5C).

The results obtained with SPX5551 were confirmed using the clinically available drug candidate balixafortide (SPX6326) (Supplementary Figure S6).

In summary, the addition of the CXCR4 inhibitor SPX5551 restores sensitivity to BTK and PI3K inhibitors in MZL models of acquired secondary resistance

### The CXCR4 inhibitor SPX5551 shows limited dose-dependent anti-tumor activity in B-cell lymphoma models as a single agent

Following the initial results in the MZL models, we took advantage of transcriptome data previously obtained in our laboratory ^38^, we selected 18 CXCR4-positive cell lines derived from mantle cell lymphoma (MCL; n=10), chronic lymphocytic leukemia (CLL; n.=3), and diffuse large B cell lymphoma (DLBCL; n.=5).

We first assessed the anti-proliferative activity of the CXCR4 inhibitor SPX5551. After 72 hours of drug exposure, the single-agent activity was minimal. Dose-dependent anti-proliferative activity was achieved in some of the MCL cell lines, while no effect was seen in the CLL or ABC DLBCL at concentrations of SPX5551 up to 100 µM (Supplementary Figure S7; Supplementary Table S3).

### Adding the CXCR4 inhibitor SPX5551 improves the anti-tumor activity of conventional and targeted therapies in B-cell lymphoma models

Based on the promising results obtained in MZL cell lines and on the literature ^9,43-46^, we tested whether adding SPX5551 could improve the anti-tumor activity of established agents. Overall, the combination of SPX5551 with conventional or targeted therapies was beneficial, additive, or synergistic across all tested B-cell lymphomas.

As observed in MZL, the combination of SPX5551 and the BTK inhibitor ibrutinib was superior to single treatments in MCL and CLL, with a stronger benefit in MCL than in CLL (Supplementary Figure S8). In the MINO or JEKO1 cell lines, SPX5551 increased the potency of ibrutinib, while in MAVER1 or Z138, adding the CXCR4 inhibitor enhanced the efficacy of the single treatment.

Combination with the PI3Kα/δ copanlisib was synergistic in MCL (Supplementary Figure S9). Adding SPX5551 mostly increased the potency of the treatment in MCL (JEKO1, SP49, Z138).

SPX5551 was combined with the CD20-targeting monoclonal antibody rituximab in the MCL and CLL models (Supplementary Figure S10). In MCL models, the combination was beneficial, showing either additivity or synergism. In particular, SPX5551 increased the efficacy (GRANTA519, JEKO1, Z138) and/or the potency (REC1, SP49, SP53, UPN1) of rituximab in MCL cell lines. The advantage of adding the CXCR4 inhibitor was minimal in CLL.

Finally, we added SPX5551 to the *in vitro* version of R-CHOP in cell lines derived from DLBCL, for which R-CHOP is still the standard regimen ^47^. The addition of CXCR4 inhibition was superior to single treatments, exhibiting additive or synergistic effects (Supplementary Figure S11). We observed an increased efficacy in OCILY19 and SUDHL2 and synergistic potency in SUDHL4.

### Combined BTK and CXCR4 inhibition increases apoptosis and inhibits NF-κB and AKT signaling than individual approaches

Considering the potential clinical development of the work, we focused on dual inhibition of CXCR4, with SPX5551, and of BTK, with ibrutinib, in MCL models.

The MCL cell lines REC1, JEKO1, and Z138 were selected for cell cycle analysis based on drug response and synergy results. In line with the observed synergy, adding SPX5551 decreased the percentage of proliferating cells, increasing the sub-G0 and apoptotic cell populations compared to the ibrutinib single agent (Figure 2).

**Figure 2.**
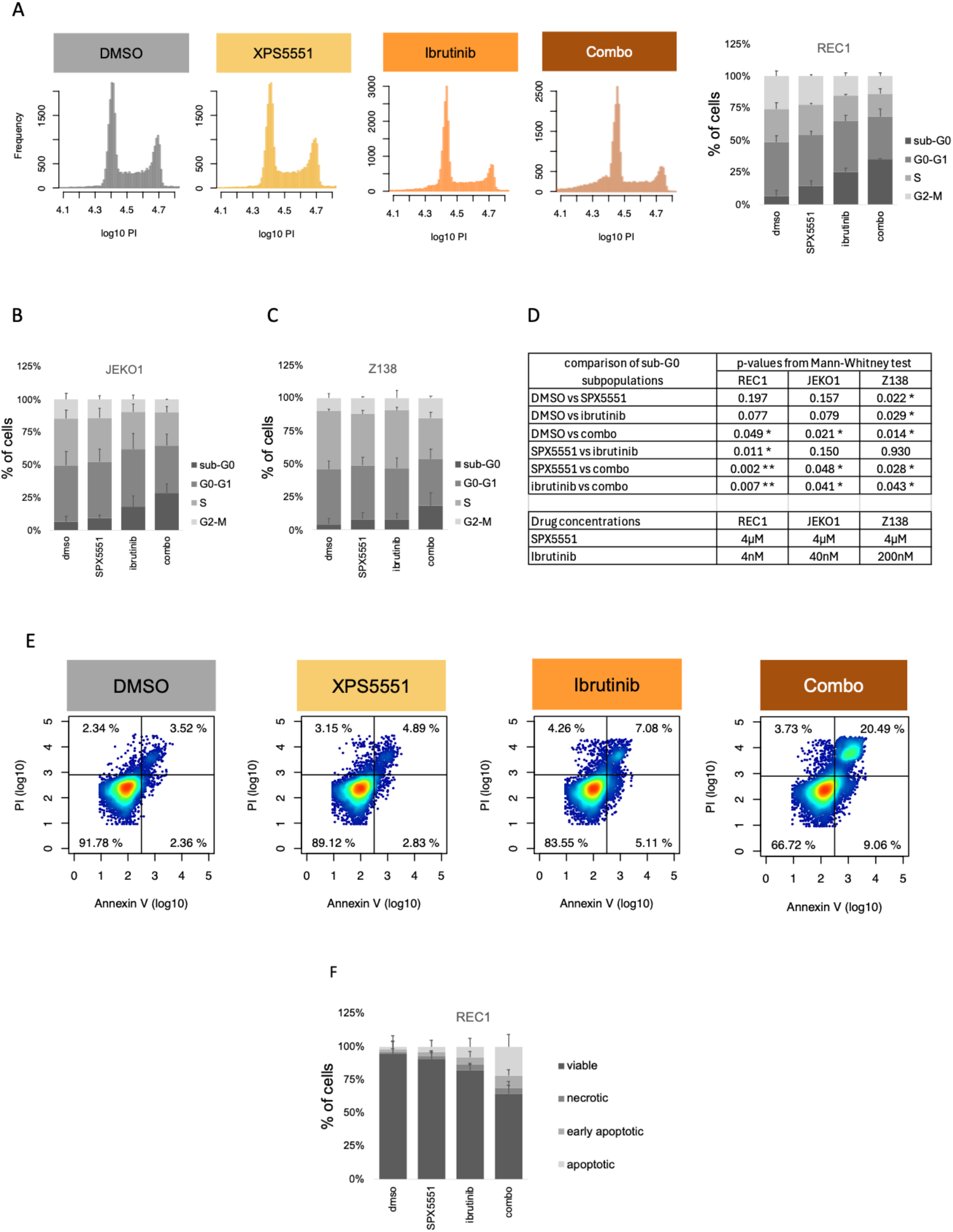
Adding SPX5551 to ibrutinib leads to increased apoptosis in MCL models. <u>Upper panel</u>, cell cycle assay upon SPX5551 and ibrutinib combination. Cell cycle phases by PI staining (FACS) obtained in REC1 (A), JEKO1 (B) and Z138 (C) models exposed to DMSO (control), SPX5551, ibrutinib, or the combination of the two. Representative barplots of two independent experiments. The table in (D) shows the corresponding p-values from the Mann-Whitney U-test comparing the sub-G0 subpopulations across treatments, * for p<0.05, ** for p>0.005. The bottom table shows the concentration used in each line. <u>Lower panel</u>, apoptosis assay by Annexin-V/PI staining obtained in REC1 cells exposed to DMSO (control), SPX5551 (4 µM), ibrutinib (4 nM), or the combination of the two. Representative scatterplots (E) and barplot (F) of two independent experiments. Apoptotic cells were significantly higher upon combo (p<0.05) but not upon any of the single agents, from a Mann-Whitney U-test comparing each treatment to DMSO.

We investigated the effect of SPX5551 and ibrutinib on NF-κB and AKT signaling. By immunofluorescence, we observed the two compounds decreased p65/RELA and p52/RELB levels, and the effect was stronger with the combination (Figure 3). The p-AKT levels were not affected by any single agent but were inhibited by the combination. As expected ^12,39^, CXCR4 was increased by ibrutinib treatment. The addition of SPX5551 counterbalanced the upregulation and repressed cytosolic CXCR4 expression (Figure 4).

**Figure 3.**
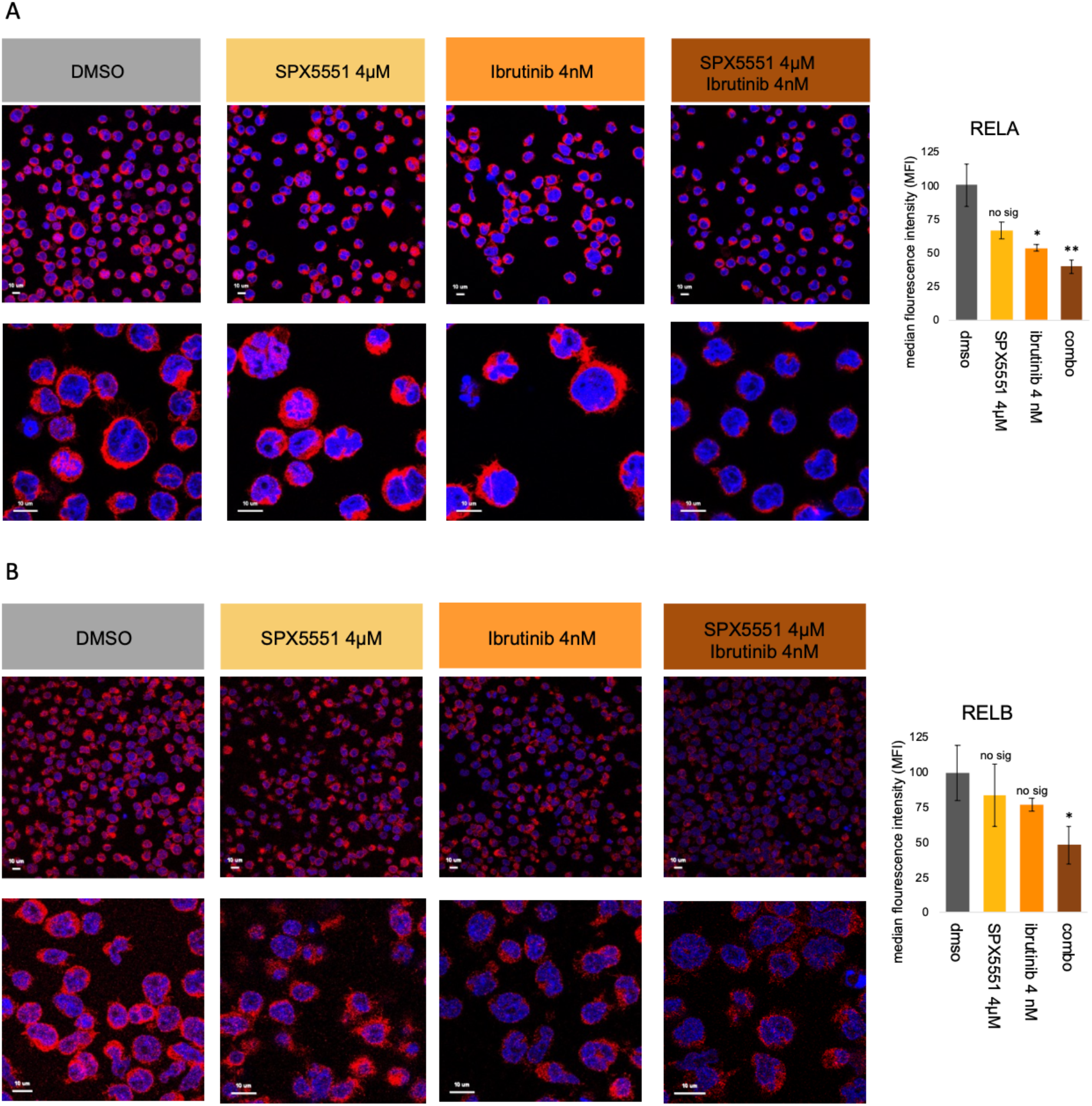
Immunofluorescence analysis of NF-κB signaling upon SPX5551 and ibrutinib combination in the MCL model REC1. REC1 cells were exposed to DMSO (control), SPX5551 (4 µM), ibrutinib (4 nM), or a combination of the two for 6 hours. Representative images of two independent experiments and protein quantification of RELA (cytoplasmatic, A) or RELB (nuclear, B). DAPI is stained blue, and RELA/RELB is stained red. Barplots on the left show the median fluorescence intensity, * for p<0.05, ** for p<0.01 from a Mann-Whitney U-test.

**Figure 4.**
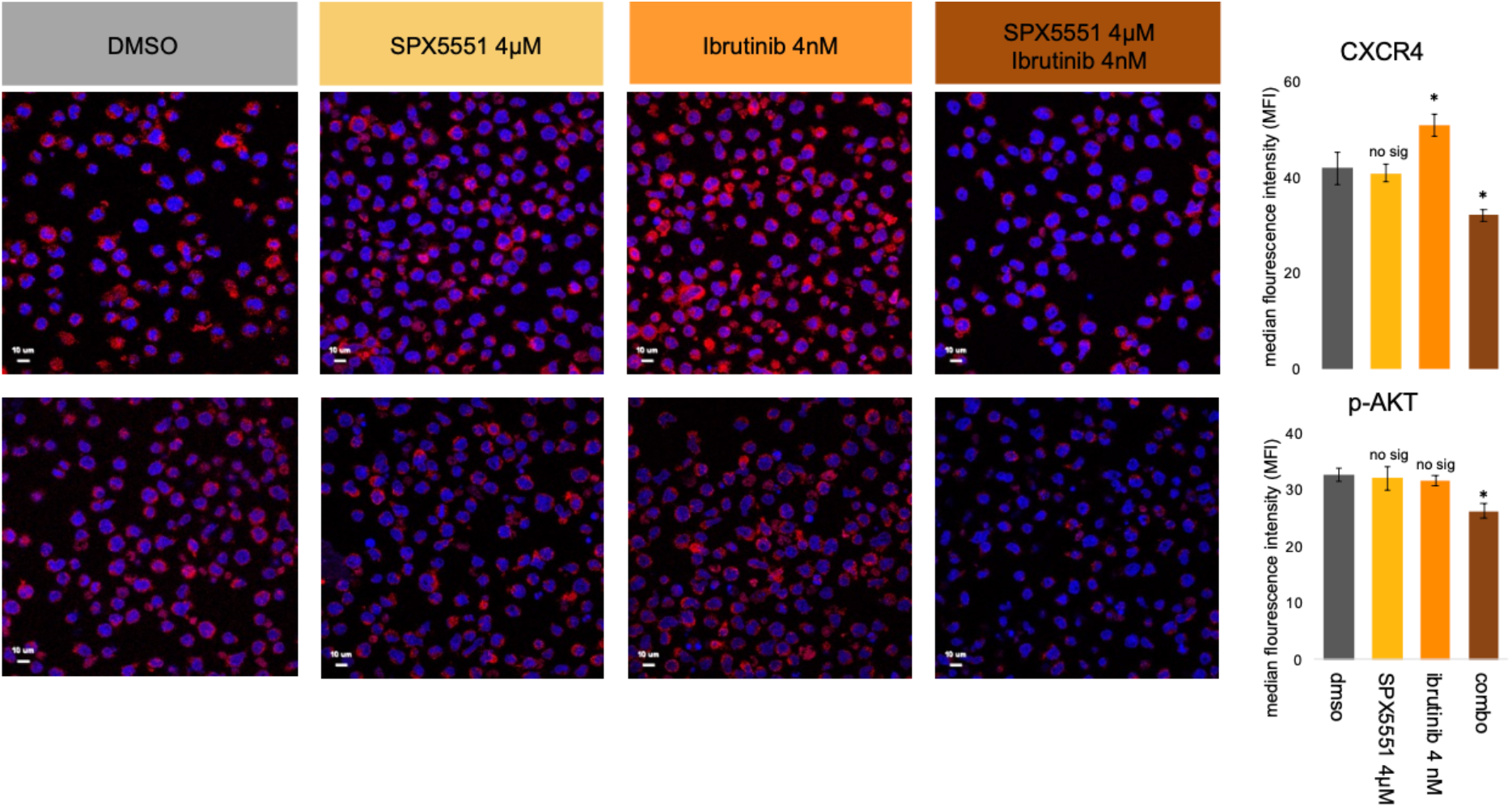
Immunofluorescence analysis of CXCR4 and p-AKT changes upon SPX5551 and ibrutinib combination in the MCL model REC1. REC1 cells were exposed to DMSO (control), SPX5551 (4 µM), ibrutinib (4 nM), or a combination of the two for 6 hours. Representative images of two independent experiments and protein quantification of CXCR4 (cytoplasmatic, top) or p-AKT (cytoplasmatic, bottom). DAPI is stained blue, and CXCR4/p-AKT is stained red. Barplots on the left show the median fluorescence intensity, * for p<0.05, ** for p<0.01 from a Mann-Whitney test.

### Combined BTK and CXCR4 inhibition has stronger effects on the transcriptome of MCL cells than individual approaches

To understand the mechanism of action of the synergy observed combining the CXCR4 and BTK inhibition, we performed transcriptome profiling in the REC1 MCL cell line exposed to SPX5551 (4 µM), ibrutinib (4 nM) or the combination of the two drugs. DMSO was used as a control. Transcriptome profiling profiles showed a much bigger impact of the SPX5551-ibrutinib combination on transcriptome than single agents: 184 transcripts differentially expressed (P <0.01, abs(log2 fold change >1) in the combination group vs DMSO, 90 (ibrutinib) and 56 (SPX5551) (Figure 5A, Supplementary Figure S12; Supplementary Table S4). Down-modulated transcripts by the combination belonged to cytokine-mediated, TNF, NF-κB, Toll-like receptor, JAK-STAT signaling pathways, and MYC targets. Compared to single agents, the combo had a stronger down-regulation of, among others, *TNF, EGR2-3, CCL4, CCL3, CXCL10, CXCR3, CD83, DUSP2, BIRC3, MIR17HG*, and *MYC* transcripts (Figure 5B-C, Supplementary Table S3). Conversely, genes up-regulated by the SPX5551-ibrutinib combination involved apoptosis, p53 and Wnt-Beta catenin signaling (Figure 5B-C, Supplementary Table S3).

**Figure 5.**
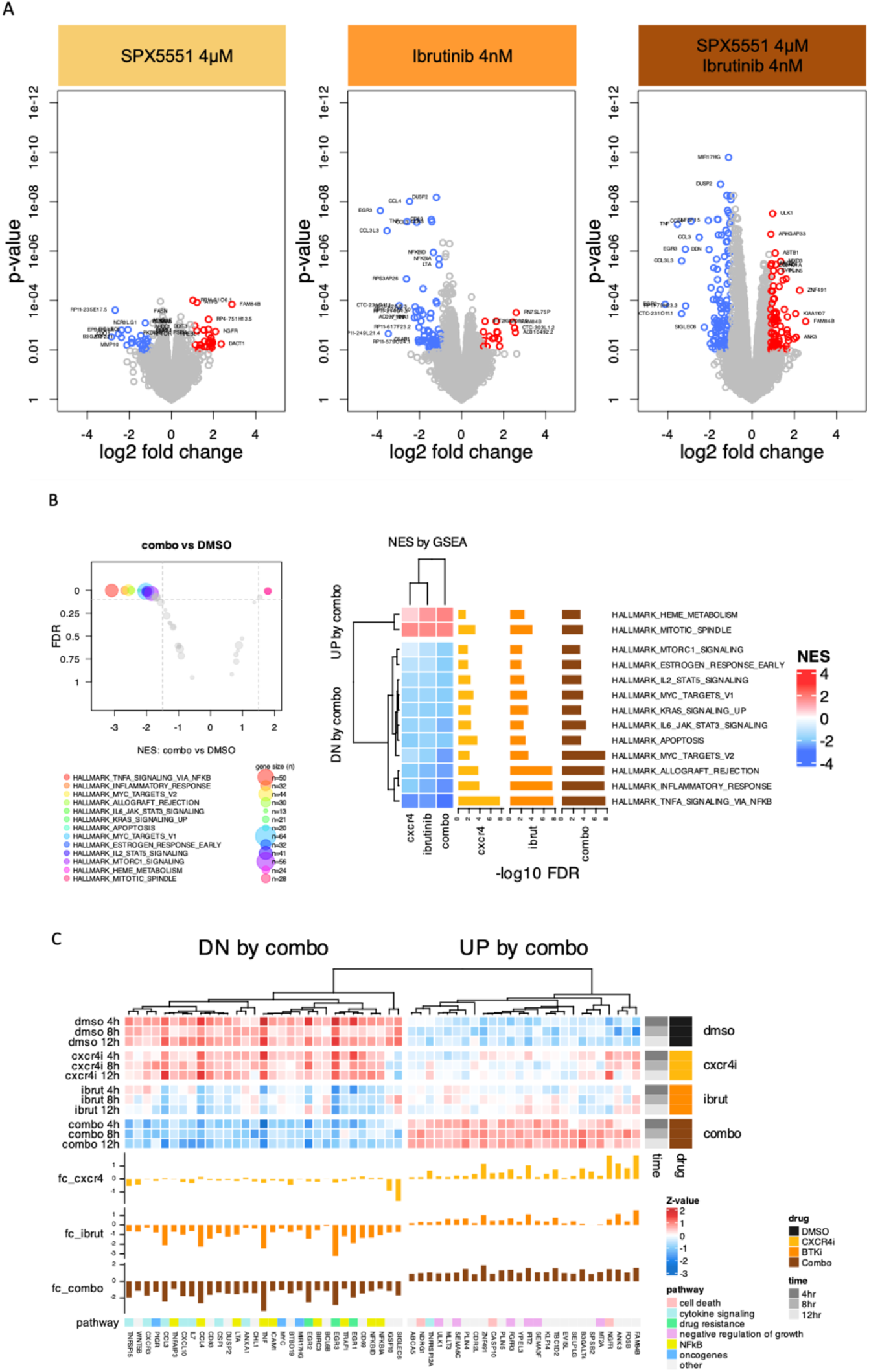
Transcriptome profiling in the MCL model REC1 cells exposed to single CXCR4 and BTK inhibitors or to the combination of the two. REC1 cells were exposed to DMSO (control), SPX5551 (4µM), ibrutinib (4nM), or the combination of the two for 4, 8 or 12 hours and poly-A RNA-sequencing was performed. (A) Volcano plots of SPX5551 vs DMSO (left), ibrutinib vs DMSO (middle) or combination vs DMSO (right), up-regulated genes in red and downregulated in blue. (B) Differentially enriched Hallmark genesets (MSigDB, Broad Institute) across treatments. FDR for false discovery rate, NES for normalized enrichment score. (C) Top overexpressed and downregulated genes by the SPX5551-ibrutinib combo in REC1 cells, fc for fold-change (log2), CXCR4i for SPX5551, BTKi for ibrutinib.

## Discussion

Here, we showed that, when combined with standard anti-lymphoma agents, CXCR4 pharmacological inhibition with the synthetic peptides SPX5551 and balixafortide enhances the overall anti-tumor effect, despite a limited single-agent activity.

Molecular simulations have been widely and successfully employed to study binding processes in CXCR4 and GPCRs in general ^48–50^. Taking advantage of the available crystallographic atomistic structure of CXCR4 in complex with a cyclic peptide ^16^ structurally similar to SPX5551 and balixafortide, we first performed a very detailed modeling showing the high similarity of the two synthetic peptides in binding to the chemokine receptor.

We then explored the anti-proliferative activity of SPX5551 as a single agent in 20 lymphoma models, which all expressed wild-type CXCR4. We observed an overt anti-tumor effect only in two cell lines. One was a cell line initially obtained from a patient with transformed splenic MZL ^51^. Since this model bears the *MYD88* L265P mutation, present in MZL but at a much lower rate than in WM/LPL ^3,52–55^, we cannot exclude that Karpas1718 represents a model of WM/LPL, a disease in which the CXCR4 can play an important role, as underlined by the frequent presence of somatic mutations in its gene ^3^. The second cell line was the MCL model SP-49. Both cell lines are also sensitive to BTK inhibitors ^56,57^, although other similarly ibrutinib-sensitive models, such as OCI-Ly10 and REC1 ^56^, were not very responsive to SPX5551. Although not reaching the IC50 with a concentration up to 100 μM, a dose-dependent activity was seen in other MCL cell lines but not in the three CLL cell lines and the five DLBCL cell lines. Our results are largely in line with what was reported using other CXCR4 antagonists ^40,44,45,58–62^, which, at low concentrations, lack a strong antitumor activity as single agents. We cannot exclude that individual molecules with specific binding modalities to CXCR4 possess a higher intrinsic ability to block the proliferation of lymphoma cells. Moreover, as we observed for Karpas1718 and SP-49, some tumor cells could depend more on CXCR4 signaling than others. A direct comparison of different molecules in a wide panel of models might help to solve this issue.

Even though the activity as a single agent was limited to very few cell lines, and in line with the notion that CXCR4 can give resistance to BCR signaling inhibitors, monoclonal antibodies, or chemotherapy ^39,40,44,58–60,63–66^, we observed a widespread benefit of adding SPX5551 to well-established anti-lymphoma agents. In MZL models, the CXCR4 pharmacological inhibition, combined with BTK or PI3K inhibitors, restored sensitivity to these drugs in cell lines derivative with acquired resistance to these agents. Among the other findings, the most interesting, also from their potential clinical translation, was that SPX5551 improved the activity of the BTK inhibitor ibrutinib, of the CD20 targeting antibody rituximab, and R-CHOP. Of interest, adding the CXCR4 inhibitor mostly resulted in increased efficacy, that is the maximal inhibitory effect of the other compound.

The benefit of adding SPX5551 or balixafortide to BTK inhibitors aligns with the notion that CXCR4 can give resistance to BCR signaling inhibitors ^39,40,63–65^ and with previous studies using different CXCR4 inhibitors ^62,66^. Importantly, the combination of BTK and CXCR4 inhibitors might be feasible in the clinics based on a phase 1 trial that explored the anti-CXCR4 antibody ulocuplumab with ibrutinib in CXCR4-mutated WM patients ^67^.

The benefit of adding SPX5551 to rituximab observed in MCL models overlaps with what was reported with plerixafor in DLBCL cell lines ^44,59^, suggesting it represents a class effect.

High CXCR4 expression has been associated with poor outcomes in DLBCL patients treated with the standard immunochemotherapy R-CHOP ^59,60^. Here, the addition of SPX5551 was beneficial when added to the *in vitro* version of the R-CHOP. This finding follows what was observed combining CXCR4 antagonists with chemotherapy agents in solid tumors ^2^.

Focusing on the combination of SPX5551 and ibrutinib, we demonstrated that it determined a stronger down-regulation of p65/RELA, p52/RELB, and p-AKT levels, and more robust down-modulation of individual transcripts, including EGR2/3, BIRC3, MIR17HG, and MYC, and pathways such as NF-κB, TLR, and JAK-STAT, leading to an increased cytotoxic activity of ibrutinib.

Based on the data we presented, it seems worth exploring CXCR4 inhibition using SPX5551 or balixafortide for patients with lymphomas mostly in combination based on the safe and well tolerated profile observed in the clinical tials performed with balixafortide as a single agent or in combination ^7,8^. Furthermore, patient selection could be important to enrich the study population of patients who could benefit from the addition of CXCR inhibition. We imagine that the liquid biopsy paired with CXCR4-targeting radio imaging or an immunohistochemistry assay in patients receiving therapies, such as BTK inhibitors, might help identify cases in which CXCR4 plays a role in resistance and could be targeted in combination with the standard therapies.

In conclusion, we have shown that the CXCR4 inhibitor SPX5551 has *in vitro* anti-tumor activity as a single agent in only a few lymphoma models, but it can increase the efficacy of targeted agents and chemotherapy. These data support further exploration of SPX5551 and its close analogue balixafortide in lymphomas.

## Author contribution

LB: performed experiments, analyzed and interpreted data, and co-wrote the manuscript.

AJA performed experiments, analyzed and interpreted data, performed data mining, prepared the figures, and co-wrote the manuscript.

LC performed data mining.

FG, FS, and CT performed experiments.

SDC, SR, VL: performed modeling computational studies.

AS, EZ, DR: provided advice.

AR performed genomics experiments;

JZ: co-designed research and provided reagents;

FB co-designed research, interpreted data, and co-wrote the manuscript. All authors reviewed and accepted the final version of the manuscript.

## Funding

This work was partially supported by institutional research funds from Spexis AG, the Swiss National Science Foundation (SNSF 31003A_163232/1) (DR, EZ, FB), Swiss Cancer Research (KFS-4727-02-2019) (AS, FB), the European Research Council (ERC) under the European Union’s Horizon 2020 research and innovation programme (“CoMMBi” ERC grant agreement No.101001784) (VL), and a grant from the Swiss National Supercomputing Centre (CSCS) under project ID u8 (VL).

## Conflict of interest

Alberto J. Arribas: travel grant from Astra Zeneca and Floratek Pharma, consultant for PentixaPharm. Luciano Cascione: institutional research funds from Orion; travel grant from HTG.

Chiara Tarantelli: travel grant from iOnctura.

Anastasios Stathis: institutional research funds from Pfizer, MSD; Roche, Novartis, Amgen, Abbvie, Bayer, ADC Therapeutics, MEI Therapeutics, Philogen, Celestia. Astra Zeneca; travel grant from AbbVie and PharmaMar; consulting fee payed to institution from Jansen, Roche, Eli Lilly.

Emanuele Zucca: institutional research funds from Celgene, Roche and Janssen; advisory board fees from Celgene, Roche, Mei Pharma, Astra Zeneca and Celltrion Healthcare; travel grants from Abbvie and Gilead; expert statements provided to Gilead, Bristol-Myers Squibb and MSD.

Davide Rossi: grant support from Gilead, AbbVie, Janssen; honoraria from Gilead, AbbVie Janssen, Roche; scientific advisory board fees from Gilead, AbbVie, Janssen, AstraZeneca, MSD.

Francesco Bertoni: institutional research funds from ADC Therapeutics, Bayer AG, BeiGene, Floratek Pharma, Helsinn, HTG Molecular Diagnostics, Ideogen AG, Idorsia Pharmaceuticals Ltd., Immagene, ImmunoGen, Menarini Ricerche, Nordic Nanovector ASA, Oncternal Therapeutics, Spexis AG; consultancy fee from BIMINI Biotech, Floratek Pharma, Helsinn, Immagene, Menarini, Vrise Therapeutics; advisory board fees to institution from Novartis; expert statements provided to HTG Molecular Diagnostics; travel grants from Amgen, Astra Zeneca, iOnctura.

The other Authors have nothing to disclose.

