## Supplementary material and methods for "Pharmacological inhibition of CXCR4 increases the anti-tumor activity of conventional and targeted therapies in B-cell lymphoma models"

*, co-contributed

**Supplementary table and figures**

**Supplementary Tables**

**Supplementary Table 1. Cell lines used in the study.**

| | | |
|---|---|---|
| **Mantle cell lymphoma** | **GRANTA-519** | Deutsche Sammlung von Mikro-organismen und Zellkulturen (DSMZ) |
| | **JEKO-1** | Eisaku Kondo, (Okayama, JP) |
| | **JVM2** | Elias Campo (Barcelona, SP) |
| | **MAVER1** | Alberto Zamò (Verona, IT) |
| | **MINO** | Robert Kridel (Vancouver, Canada) |
| | **REC-1** | Finbarr Cotter (London, UK) |
| | **SP49** | Robert Kridel (Vancouver, Canada) |
| | **SP53** | Robert Kridel (Vancouver, Canada) |
| | **UPN-1** | Robert Kridel (Vancouver, Canada) |
| | **Z-138** | Robert Kridel (Vancouver, Canada) |
| **Chronic lymphocytic leukemia** | **MEC1** | Alberto Zamò (Verona, IT) |
| | **PCL12** | Cristina Scielzo (Milan, IT) |
| | **HG3** | Deutsche Sammlung von Mikro-organismen und Zellkulturen (DSMZ) |
| **Diffuse large B-cell lymphoma** | **OCI-Ly10** | Laura Pasqualucci (New York, NY, USA) |
| | **OCI-Ly19** | Louis M. Staudt (Bethesda, MD, USA) |
| | **U2932** | Bettina Borisch (Geneva, CH) |
| | **SU-DHL-2** | Laura Pasqualucci (New York, NY, USA) |
| | **SU-DHL-4** | Laura Pasqualucci (New York, NY, USA) |
| **Marginal zone lymphoma** | **Karpas1718** | José Ángel Martinez-Climent (Pamplona, SP) |
| | **VL51** | José Ángel Martinez-Climent (Pamplona, SP) |

**Supplementary Table S2. Concentrations of drugs used in each cell line.**

**Supplementary Table S3. IC50 and combination data.**

**Supplementary Table S4. Limma and GSEA.**

### Supplementary Figures

**Supplementary Figure 1**.
Scatterplot showing the convergence in the Rosetta Docking calculations for balixafortide and SPX5551 (left and right panels, respectively). On the x-axis, we report the RMSD of the cyclic peptides' $C_\alpha$ atoms computed with respect to the top-scoring binding mode, labeled with a star. Each value is calculated after aligning the Cα atoms of the CXCR4 binding pocket's residues within 4 Å of the ligand. On the y axis, we report the score of each pose as Rosetta Energy Units (REU), computed through the ref2015 score function of Rosetta docking software. Furthermore, we show the RMSD value computed for the $C_\alpha$ atoms of the crystal structure of CVX15 with respect to the top-scoring binding mode (red lines).

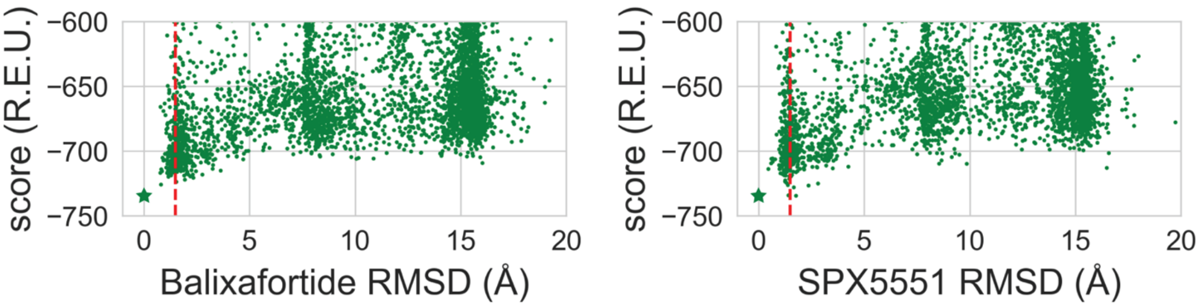

**Supplementary Figure 2.** A) Structural comparison of the binding modes of balixafortide, CVX15 and SPX5551 to CXCR4. Three-dimensional representation of the binding modes of balixafortide, CVX15 and SPX5551 to CXCR4. The protein is represented as grey ribbons, the docked peptides as green ribbons, and CVX15 as tan ribbons. The interacting residues are displayed as sticks, and the interactions are shown as dashed black lines. In detail, all three peptides assume a disulfide-stabilized (Cys4 to Cys11) β-hairpin, with D-Pro15 and Pro16 at the tip of the turn exposed to the extracellular milieu. The first difference between CVX15 and the two docked peptides is at residue 2. In this position, CVX15 has an arginine instead of a histidine in balixafortide and SPX5551. This difference impacts the type of interaction established with Phe189 of CXCR4. Specifically, in the case of CVX15, a cation-π interaction takes place between Arg2 and Phe189, whereas in balixafortide and SPX5551, His2 engages a π-π stacking interaction with Phe189. However, cation-π interaction might also occur between Phe189 and His2, depending on the protonation state of the latter. In CVX15, Arg8 establishes salt bridge interactions with Asp187 (ECL2) and Glu32 (N-terminal). Similarly, in balixafortide and SPX5551, DAB8, which replaces Arg8, forms ionic interactions with Asp97 (TM2) and Asp187 (ECL2). Arg9, which is conserved in all three peptides, forms H-bond interactions with Thr117, whereas Tyr10 in balixafortide and SPX5551 forms H-bonds with Tyr255. This interaction is lost in CVX15, where Tyr10 is replaced by Ala10. B) Multiple sequence alignment of CVX15, SPX5551, and balixafortide. Multiple sequence alignment is obtained using the Clustal Omega algorithm(1). Identical residues are displayed as white letters in red boxes, while conservative mutations are shown as red letters. Residues in D configuration are noted with an asterisk; non-canonical residues are flagged with a dagger (i.e.: $R^{\dagger}$ corresponds to citrulline, $X^{\dagger}$ to the 2-4-diaminobutyric acid, and $A^{\dagger}$ to naphtalen-2-yl-3-alanine). CVX15 numbering has been changed with respect to the PDBID 3OE0 to match the numbering used for balixafortide and SPX5551.

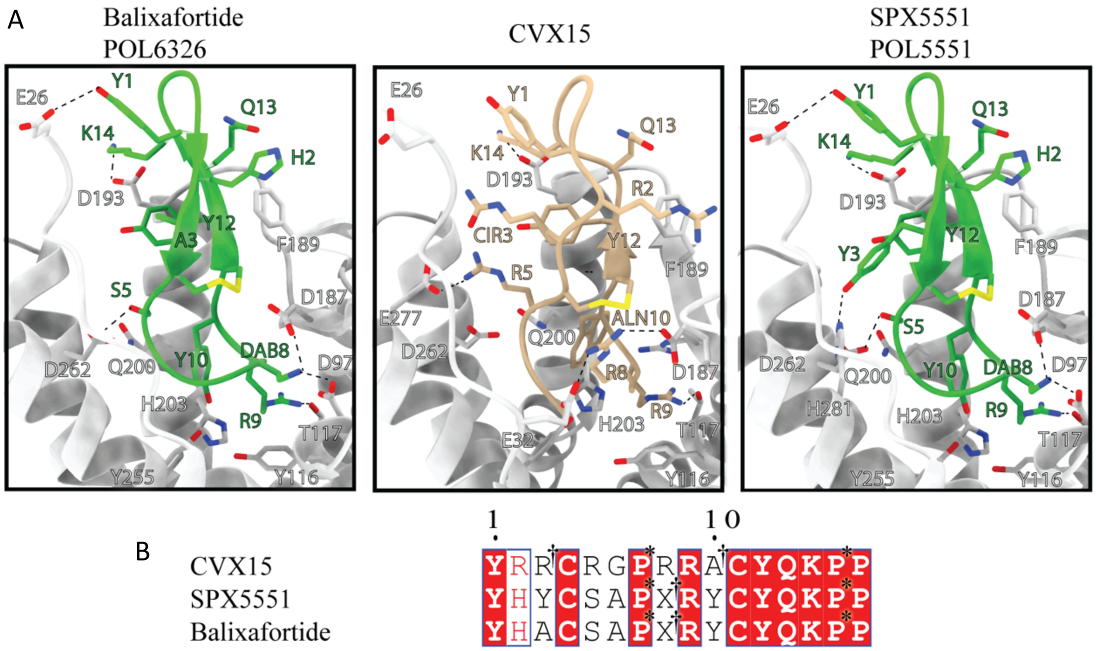

**Supplementary Figure 3.** Surface CXCR4 expression was measured by flow cytometry in VL51 and Karpas1718 parental and resistant lines.

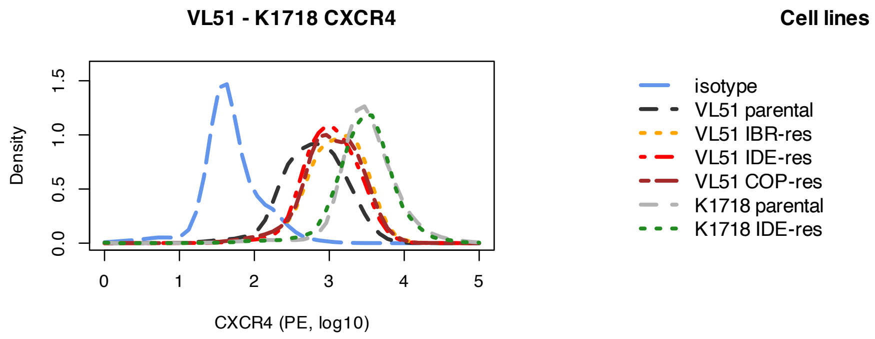


**Supplementary Figure 4.** Single-agent activity of the CXCR4 inhibitors SPX5551 and Balixafortide in MZL models with acquired resistance. Drug response curves of SPX5551 (A-B) or Balixafortide (C-D) in MZL models with acquired resistance to PI3K, BTK, or PI3K/BCL2 inhibitors (A and C derived from VL51, B and D derived from Karpas1718, K1718). Viability was assessed by MTT assay after 72 hours of exposure to increasing doses of SPX5551. Curves correspond to the mean of three independent experiments. Error bars represent the standard deviation of the mean. Supp Table 1 shows the IC50 values of SPX5551 and balixafortide upon 72 hours of exposure (4-parameter log-regression equation, "drc" package in R environment).

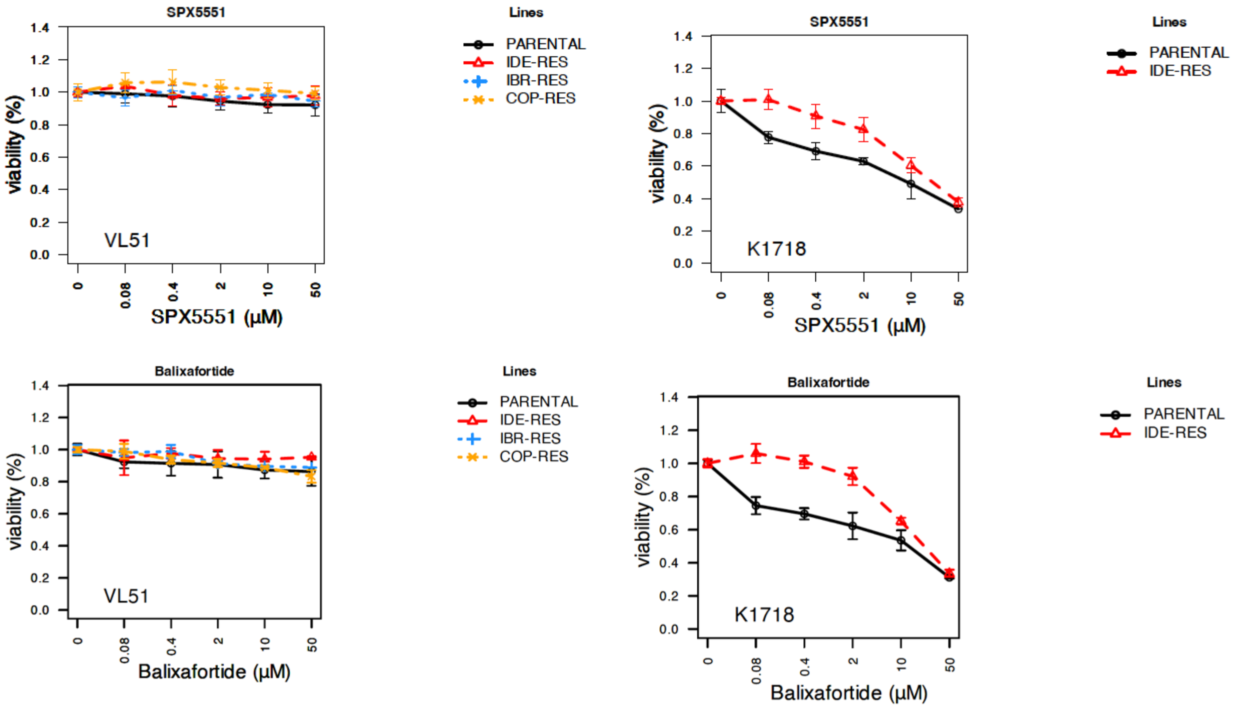

**Supplementary Figure 5.** Combination data adding SPX5551 to Ibrutinib, idelalisib and copanlisib in MZL cell lines. Combination of SPX5551 with the BTK inhibitor ibrutinib (A), and with the PI3K inhibitors Idelalisib (B) or Copanlisib (C) in VL51 (upper panel) and Karpas1718 (lower panel) cells and their derivatives with acquired resistance to the corresponding compound. Cell viability was assessed by MTT assay after 72 hours of exposure. Heatmaps show the mean of three independent experiments. The benefit of the combination was assessed both as synergism according to the Chou-Talalay combination index (left, CI: synergistic CI<0.9, additive CI~1, antagonistic CI>1) (2) and as efficacy (center, synergistic: efficacy>1, additive: 0<efficacy<1, antagonistic: efficacy<0) and potency (right, synergistic: potency>0.5, additive: 0<potency<0.5, antagonistic: potency<0) according to the MuSyC algorithm (3). (D) The addition of SPX5551 (1 μM) overcomes resistance to BTK or PI3K inhibitors in VL51 (upper panel) and Karpas1718 (lower panel) resistant derivatives.

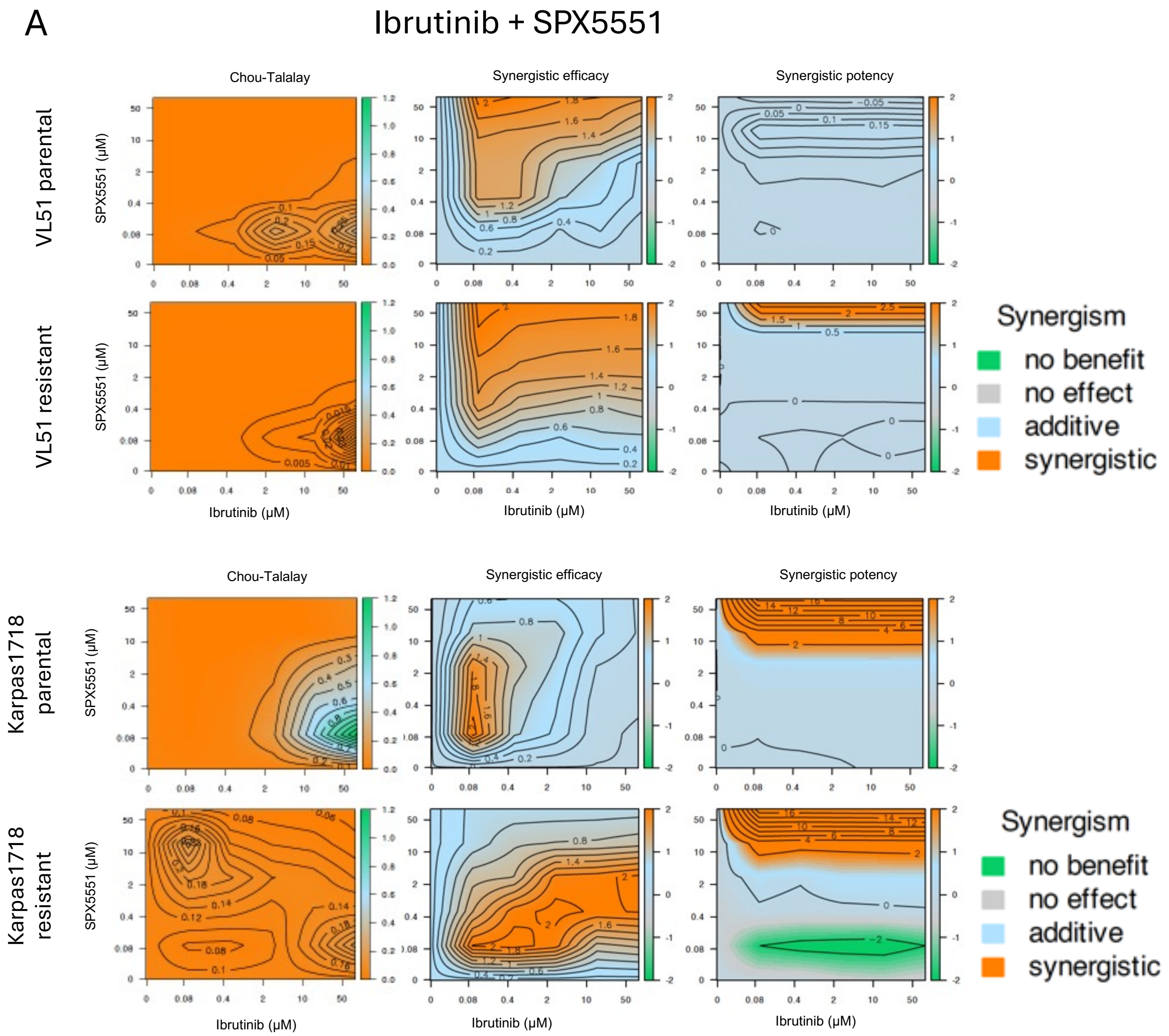

B

### Idelalisib + SPX5551

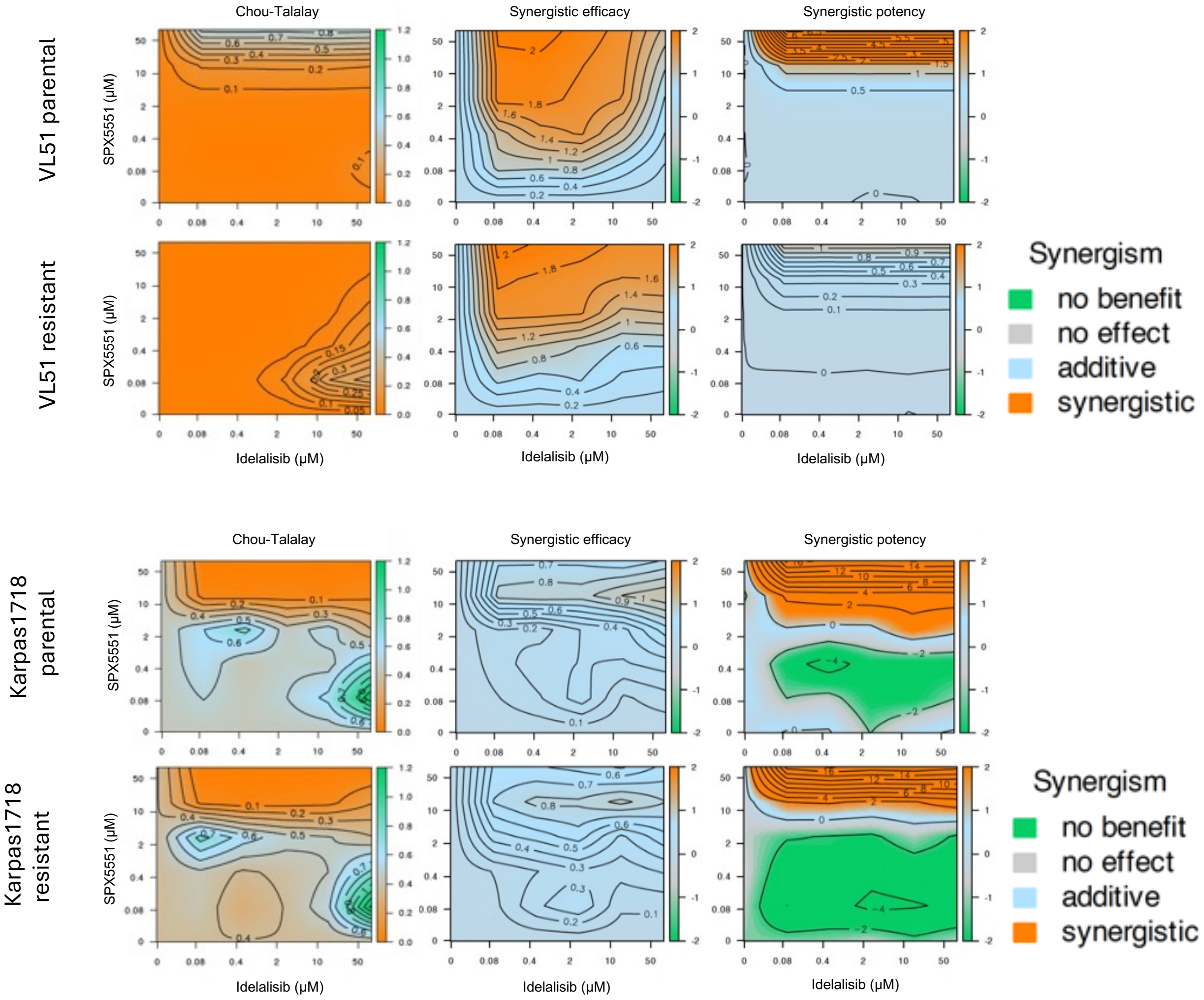

C Copanlisib + SPX5551

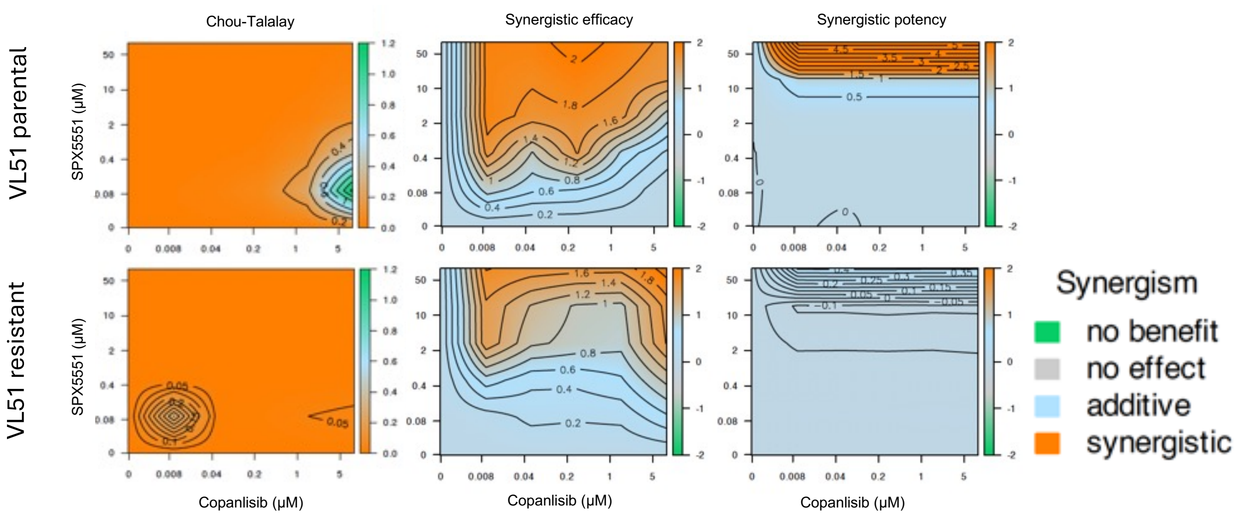

D SPX5551 restored sensitivity in resistant lines

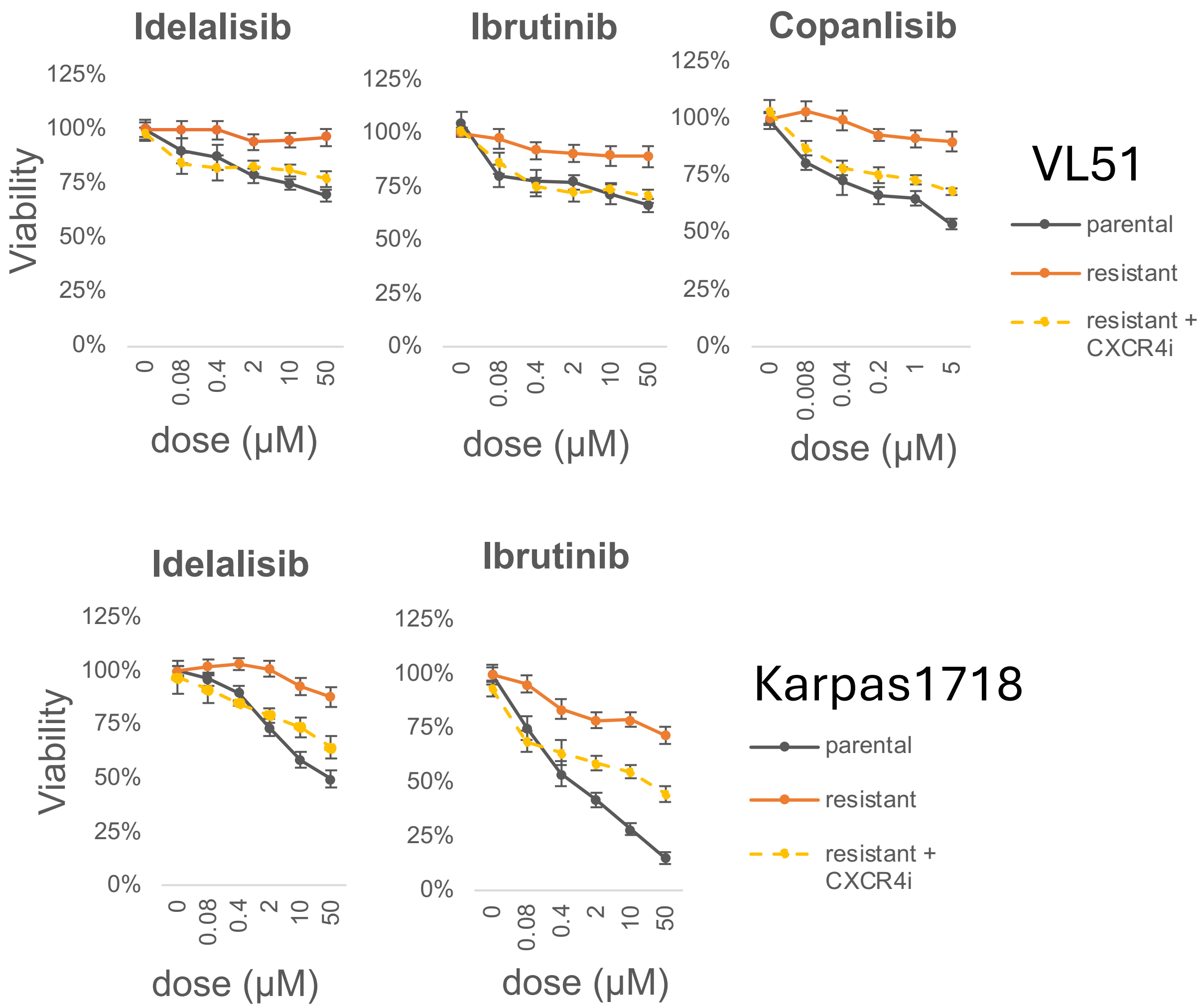

**Supplementary Figure 6.** Combination of Balixafortide with the BTK inhibitor ibrutinib (A), and with the PI3K inhibitors Idelalisib (B) or Copanlisib (C) in VL51 and Karpas1718 cells and their derivatives with acquired resistance to the corresponding compound. Cell viability was assessed by MTT assay upon 72 hours of exposure. Heatmaps show the mean of three independent experiments. The benefit of the combination was assessed both as synergism according to the Chou-Talalay combination index (left, CI: synergistic CI<0.9, additive CI~1, antagonistic CI>1) (2) and as efficacy (center, synergistic: efficacy>1, additive: 0<efficacy<1, antagonistic: efficacy<0) and potency (right, synergistic: potency>0.5, additive: 0<potency<0.5, antagonistic: potency<0) according to the MuSyC algorithm (3).

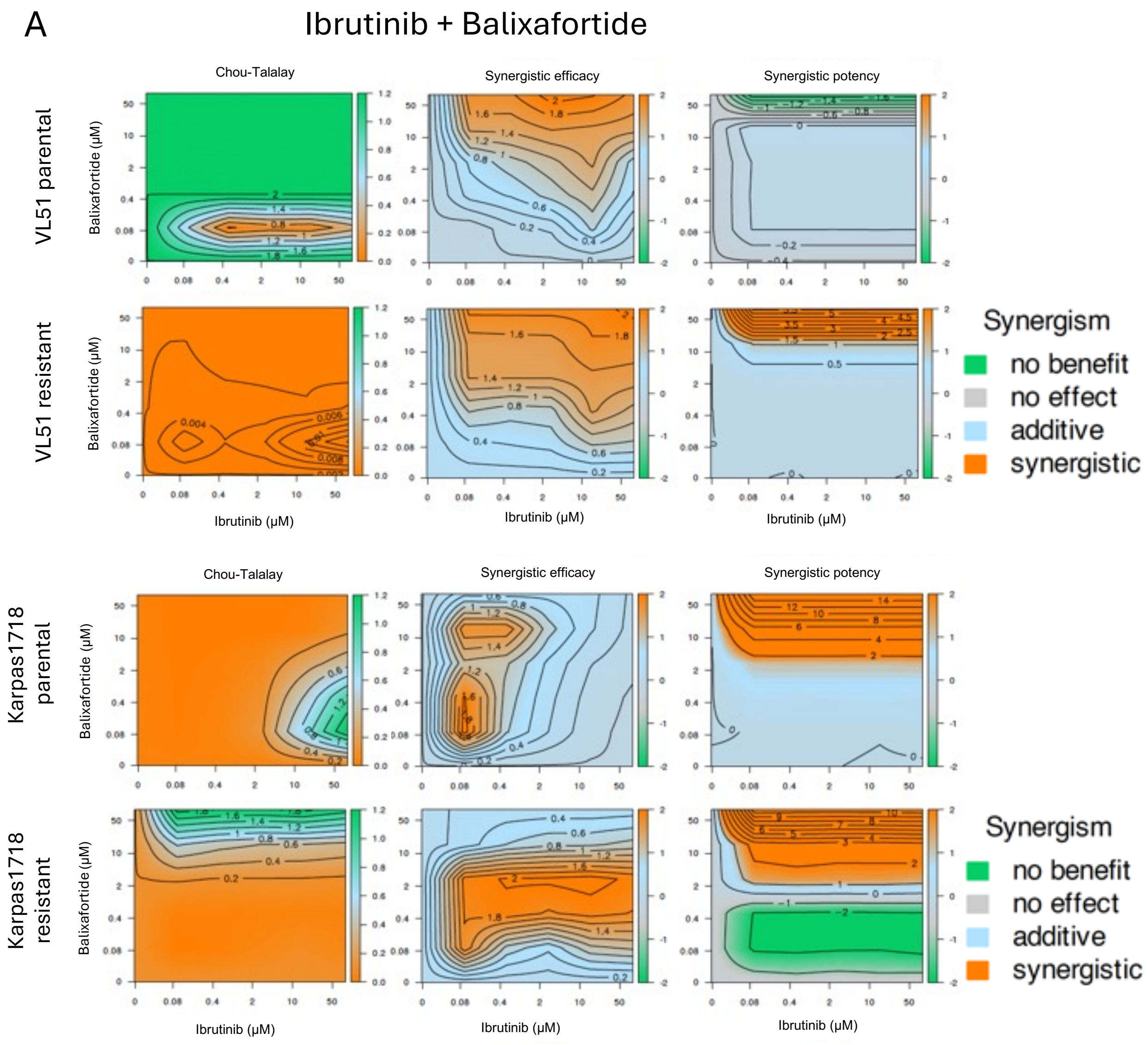

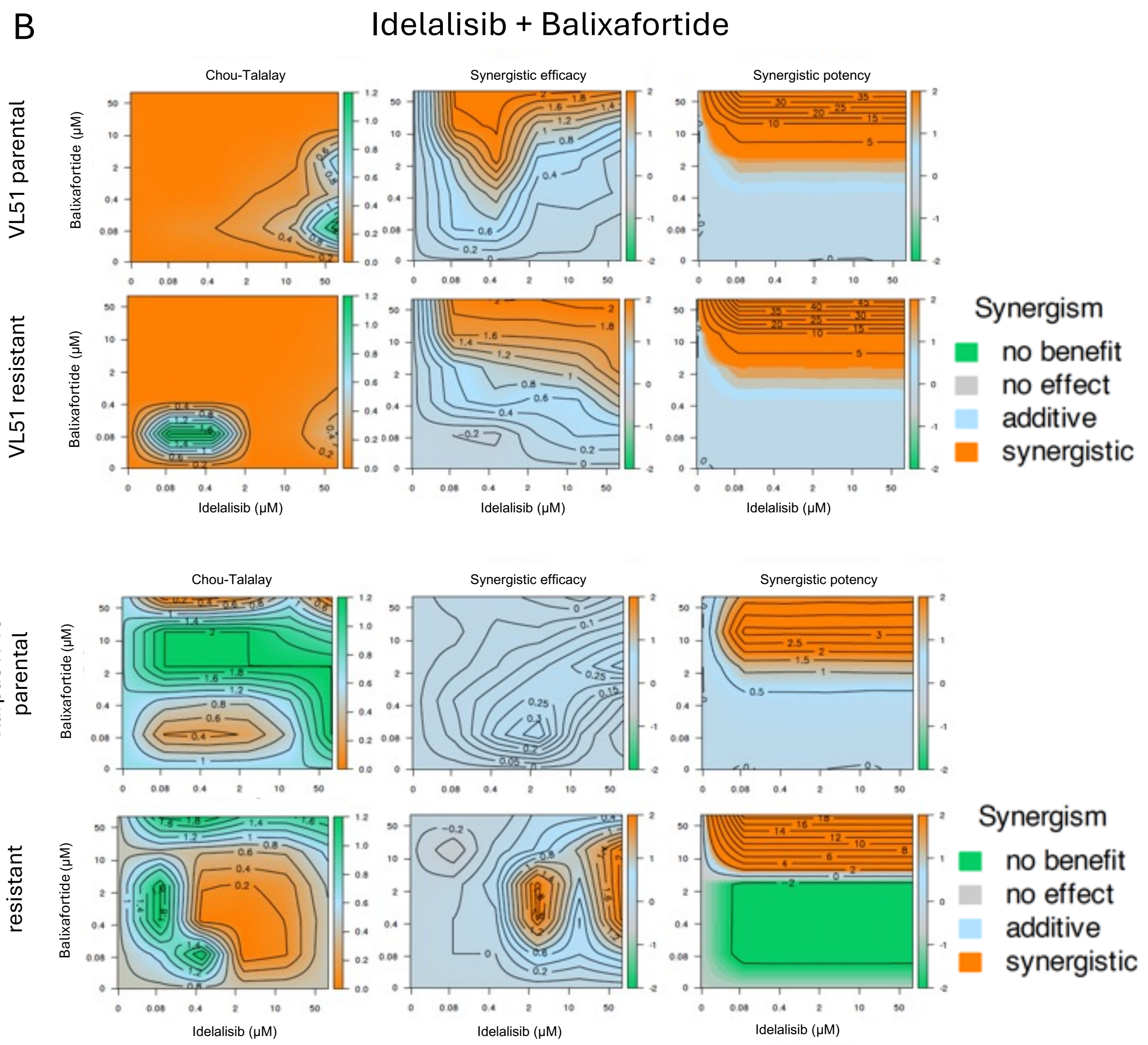
B
Idelalisib + Balixafortide
Chou-Talalay
Synergistic efficacy
Synergistic potency
VL51 parental
VL51 resistant
Karpas1718 parental
Karpas1718 resistant
Balixafortide (μM)
Idelalisib (μM)
Synergism
no benefit
no effect
additive
synergistic

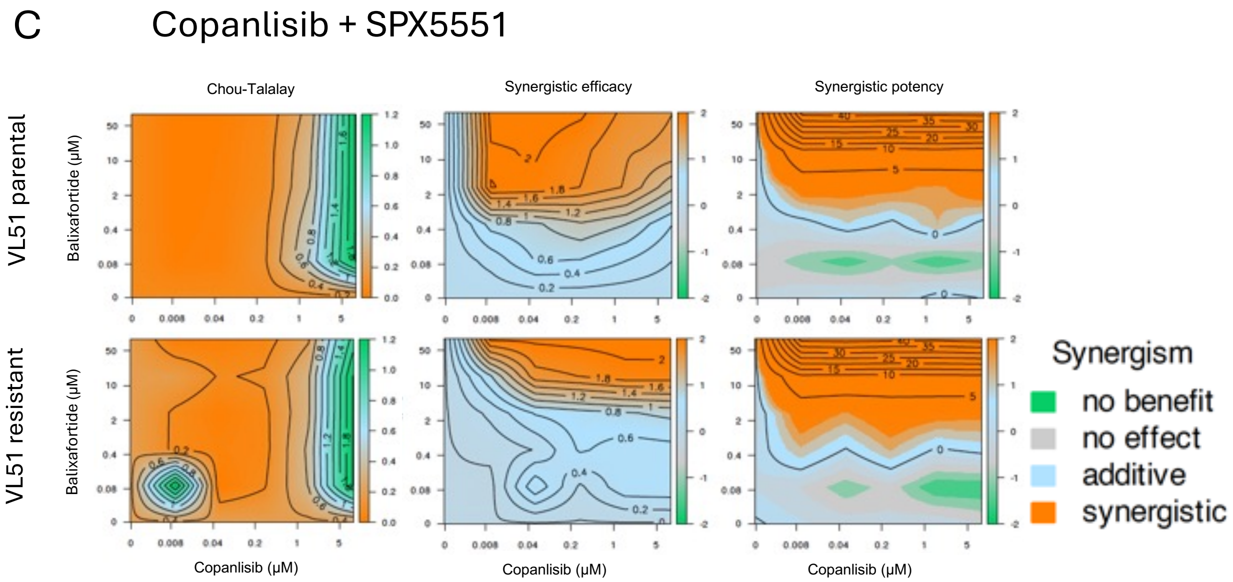
C
Copanlisib + SPX5551
Chou-Talalay
Synergistic efficacy
Synergistic potency
VL51 parental
VL51 resistant
Balixafortide (µM)
Copanlisib (µM)
Synergism
no benefit
no effect
additive
synergistic

**Supplementary Figure 7.** Single-agent activity of CXCR4 inhibitor SPX5551. Drug response curves of SPX5551 in MCL (A), CLL (B), or DLBCL (C) cell lines. Viability was assessed by MTT assay after 72 hours of exposure to increasing doses of SPX5551. Curves correspond to the mean of three independent experiments. Error bars represent the standard deviation of the mean. (D) IC50 values of SPX5551 upon 72 hours of exposure were estimated by the 4-parameter log-regression equation (“drc” package in R environment).

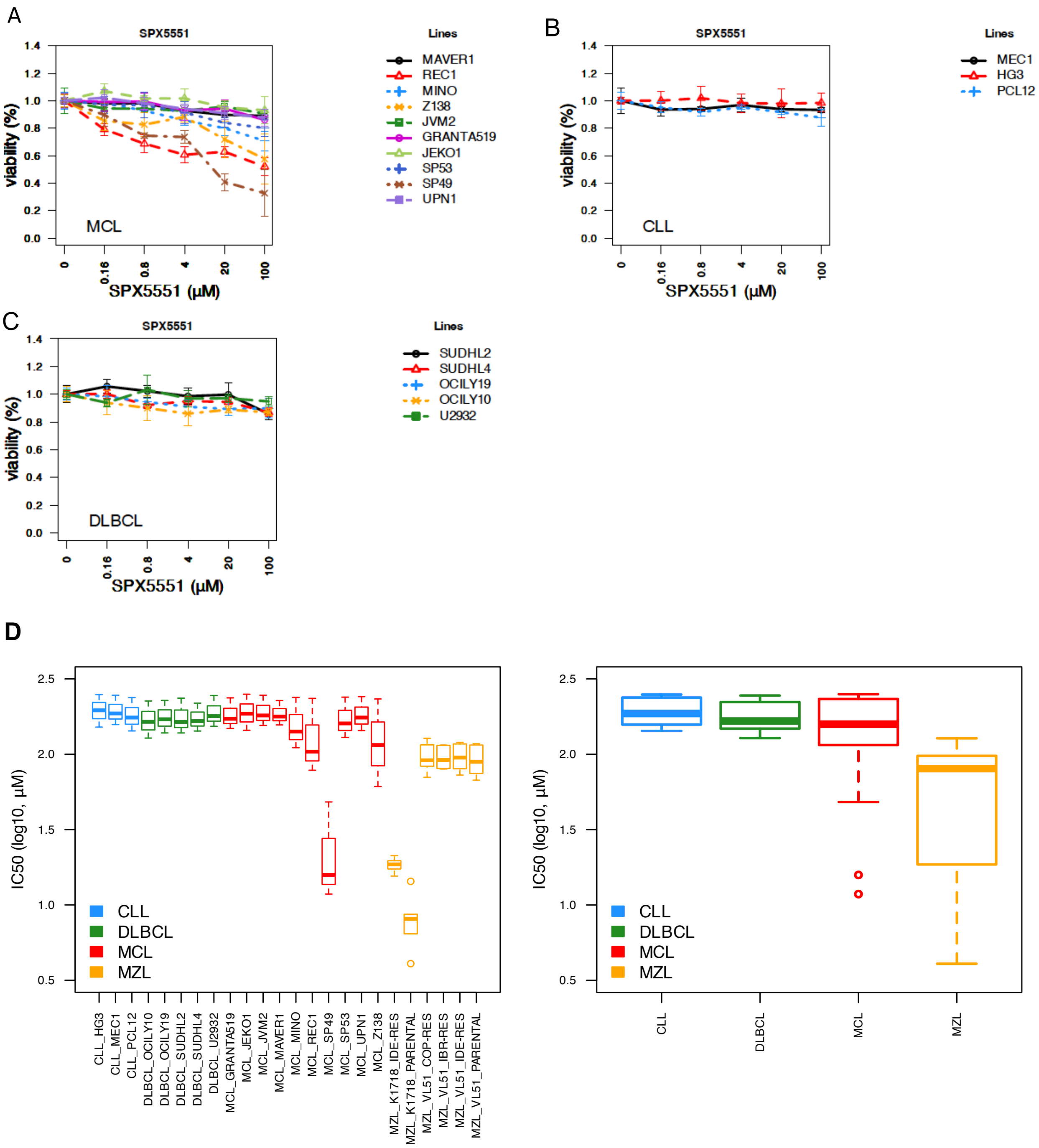

**Supplementary Figure 8**. Combination of SPX5551 with the BTK inhibitor ibrutinib in MCL (n.=10, red) or CLL (n.=3, blue) cell lines. Cell viability was assessed by MTT assay after 72 hours of exposure. Boxplots show the mean of three independent experiments. The benefit of the combination was assessed both as synergism according to the Chou-Talalay combination index (CI, synergistic: CI<0.9, additive: CI~1, antagonistic: CI>1) (A) (2) and as efficacy (synergistic: efficacy>1, additive: 0<efficacy<1, antagonistic: efficacy<0) (B) and potency (synergistic: potency>0.5, additive: 0<potency<0.5, antagonistic: potency<0) (C) according to the MuSyC algorithm (3).

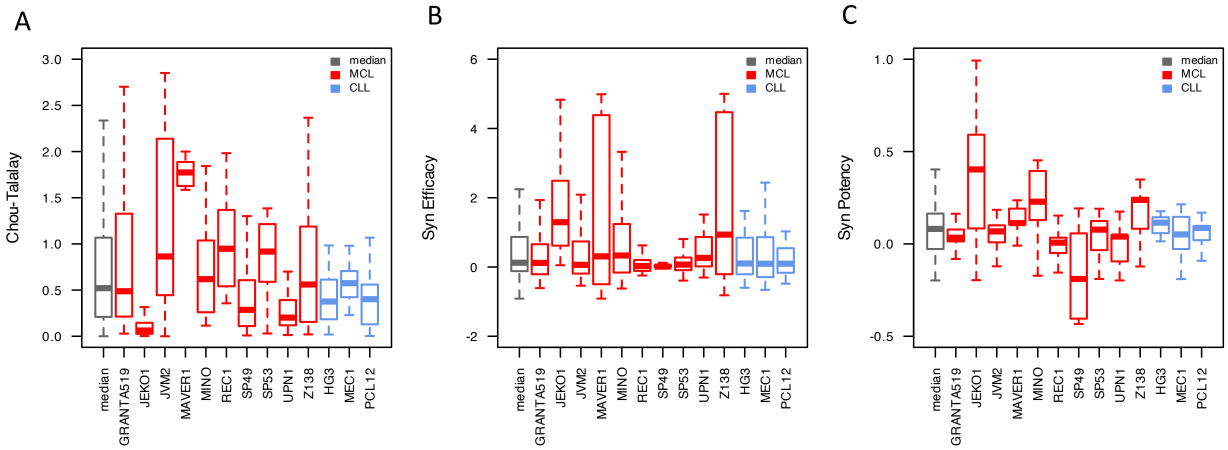

**Supplementary Figure 9.** Combination of SPX5551 with the PI3K inhibitor copanlisib in MCL (n.=10, red) cell lines. Cell viability was assessed by MTT assay after 72 hours of exposure. Boxplots show the mean of three independent experiments. The benefit of the combination was assessed both as synergism according to the Chou-Talalay combination index (CI, synergistic: CI<0.9, additive: CI~1, antagonistic: CI>1) (A) (2) and as efficacy (synergistic: efficacy>1, additive: 0<efficacy<1, antagonistic: efficacy<0) (B) and potency (synergistic: potency>0.5, additive: 0<potency<0.5, antagonistic: potency<0) (C) according to the MuSyC algorithm (3).

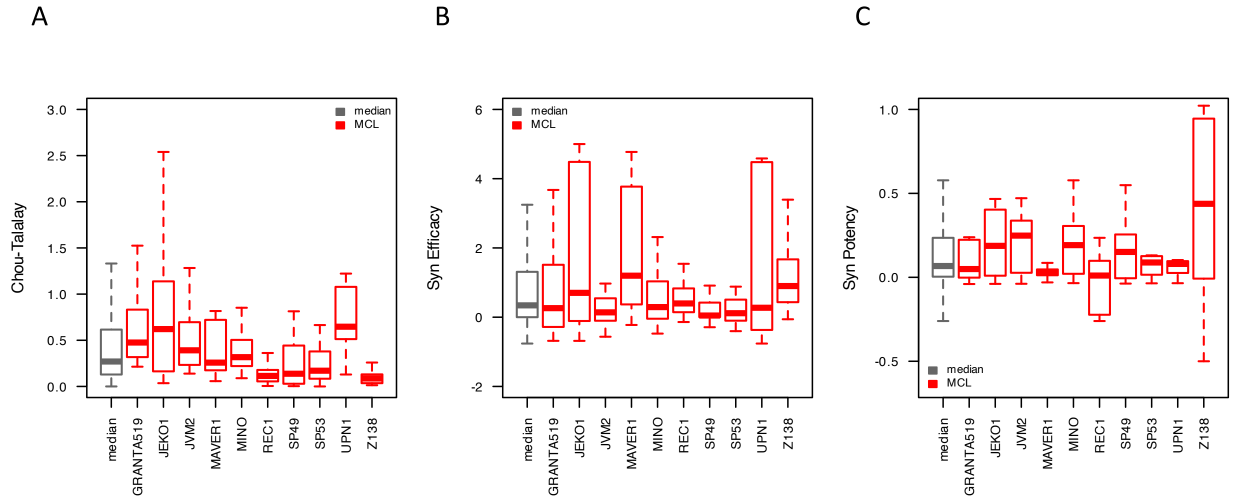

**Supplementary Figure 10.** Combination of SPX5551 with the anti-CD20 monoclonal antibody rituximab in MCL (n.=10, red) or CLL (n.=3, blue) cell lines. Cell viability was assessed by MTT assay after 72 hours of exposure. Boxplots show the mean of three independent experiments. The benefit of the combination was assessed both as synergism according to the Chou-Talalay combination index (CI, synergistic: CI<0.9, additive: CI~1, antagonistic: CI>1) (A) (2) and as efficacy (synergistic: efficacy>1, additive: 0<efficacy<1, antagonistic: efficacy<0) (B) and potency (synergistic: potency>0.5, additive: 0<potency<0.5, antagonistic: potency<0) (C) according to the MuSyC algorithm (3).

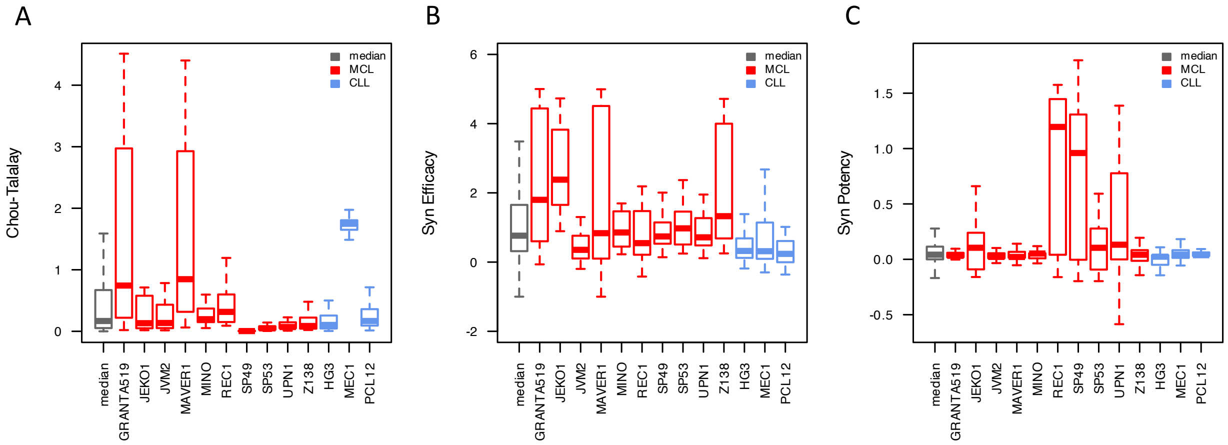

**Supplementary Figure 11.** Combination of SPX5551 with the *in vitro* version of R-CHOP in DLBCL (n.=5, green). Cell viability was assessed by MTT assay after 72 hours of exposure. Boxplots show the mean of three independent experiments. The benefit of the combination was assessed both as synergism according to the Chou-Talalay combination index (CI, synergistic: CI<0.9, additive: CI~1, antagonistic: CI>1) (A) (2) and as efficacy (synergistic: efficacy>1, additive: 0<efficacy<1, antagonistic: efficacy<0) (B) and potency (synergistic: potency>0.5, additive: 0<potency<0.5, antagonistic: potency<0) (C) according to the MuSyC algorithm (3).

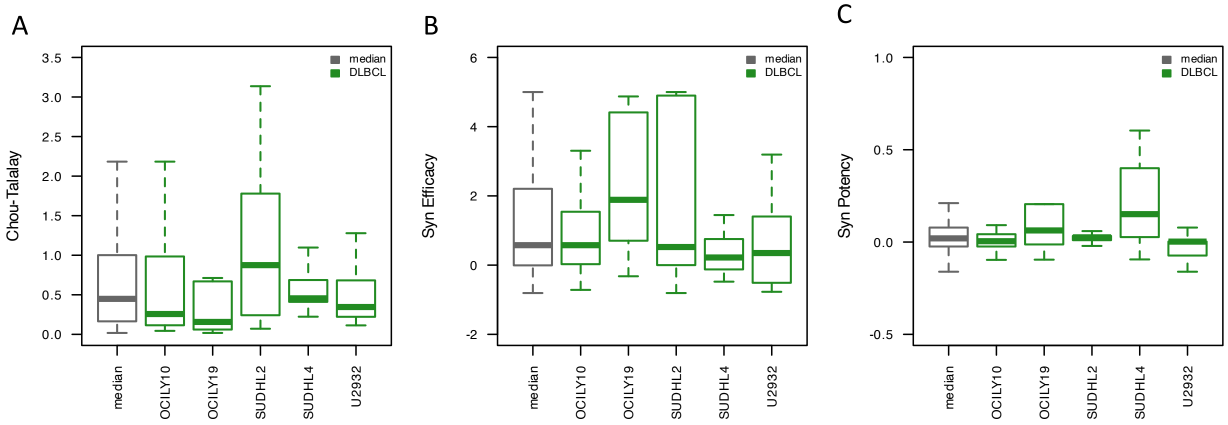

**Supplementary Figure 12.** Barplot showing the differentially expressed genes (p-value >0.01, absolute log-2 fold change >1), up (red), down (blue), or total (grey) across treatments compared to DMSO.

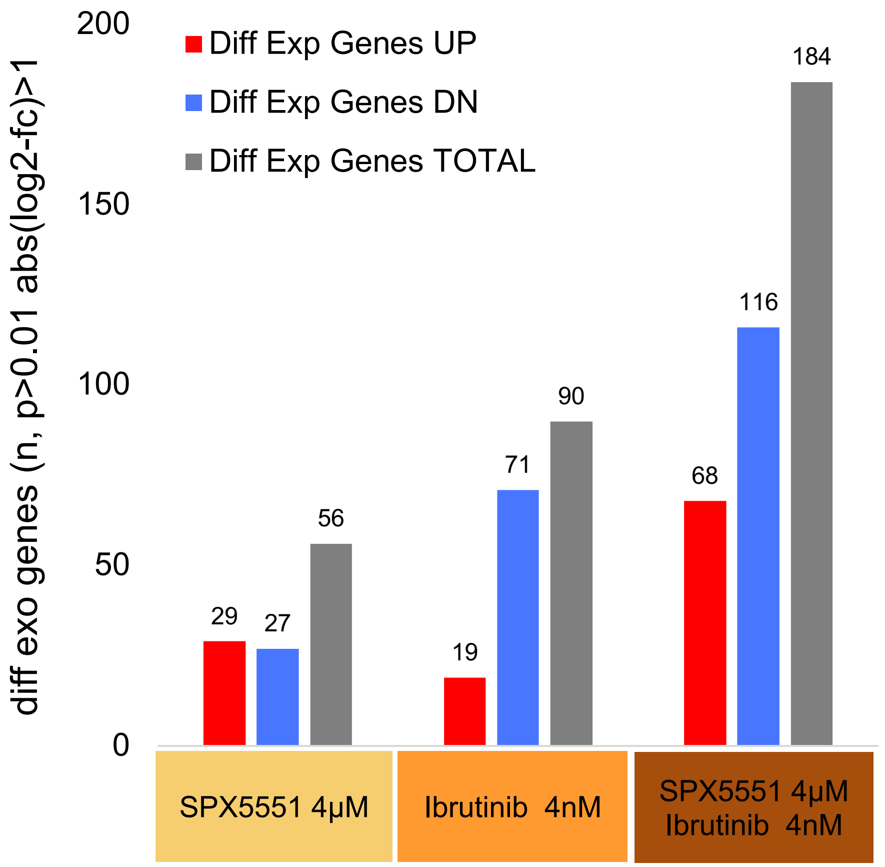
